# Engineering plants tolerant to the toxic proline mimic azetidine-2-carboxylic acid through co-option of a bacterial detoxification mechanism

**DOI:** 10.64898/2026.09.25.754441

**Authors:** Varun Dwivedi, Meghan Boozer, Craig A. Schenck

**Author notes:** Corresponding author –.

## Abstract

Azetidine-2-carboxylic acid (Aze) is a toxic, non-proteogenic amino acid produced in restricted plants. It can enter the human food chain through direct or indirect dietary exposure and has been linked to neurological disorders due to its ability to misincorporate for its structural analog proline (Pro) during translation and lead to protein misfolding. Although several microorganisms have evolved mechanisms to detoxify Aze, these pathways have not previously been exploited to engineer Aze tolerance in plants. Here, we heterologously expressed a bacterial Aze hydrolase (HAS) in *Arabidopsis thaliana* to establish a metabolic strategy for Aze detoxification in plants. Transgenic plants exhibited markedly enhanced tolerance to Aze, as demonstrated by improved root growth, cotyledon development, and survival compared with wild-type plants under Aze stress. We further showed that HAS localized to the cytosol, where it converted Aze into the non-toxic metabolite 2-hydroxy-4-aminobutyrate (HAB). HAB formation was confirmed both *in planta* and *in vitro*, demonstrating that HAS remained catalytically active in plant cells and effectively detoxified Aze. Enzyme kinetics revealed that HAS has a high affinity for Aze and is stereospecific for L-Aze, with no activity toward D-Aze. Multi-omics analysis including transcriptomics and misincorporation proteomics show that the transgenic lines misincorporate Aze significantly less compared to wild-type leading to enhanced Aze tolerance. Our findings establish bacterial HAS as an effective metabolic detoxification system that confers enhanced Aze tolerance in plants. This study provides a promising strategy for engineering plant resistance to toxic non-proteogenic amino acids.

## Introduction

Plants produce a vast array of chemically diverse specialized metabolites that mediate ecological interactions and influence organisms across all domains of life. Among these compounds are non-proteogenic amino acids, which can function as defense metabolites. Despite not being genetically encoded for protein synthesis, non-proteogenic amino acids can interfere with protein biosynthesis due to their structural similarity to proteogenic amino acids (Jander et al. 2020; Demash et al. 2025). Azetidine-2-carboxylic acid (Aze) is a naturally occurring non-proteogenic amino acid produced by several plant species that deters herbivores and restricts microbial pathogens (Fowden 1955; Huang et al. 2011; Han et al. 2022; Gibson et al. 2025). Aze has also been proposed as a potential environmental risk factor for neurodegenerative disorders, including amyotrophic lateral sclerosis (ALS), multiple sclerosis (MS), and Parkinson’s disease (Rubenstein 2008; Rodgers 2014; Rodgers et al. 2025; Sobel et al. 2026). Human exposure to Aze can occur directly through the consumption of Aze-containing crops (including beets) or indirectly from animals fed Aze-containing crops (Rodgers et al. 2025). Aze toxicity arises primarily from its structural similarity to proline (Pro), which allows it to interfere with protein biosynthesis and exert proteotoxic effects (Thives Santos et al. 2024; Demash et al. 2025). Unlike Pro, Aze contains a rigid four-membered azetidine ring that imposes distinct conformational constraints when incorporated into proteins, thereby destabilizing protein structure, promoting misfolding, and impairing function (Bessonov et al. 2010; Song et al. 2017; Rodgers et al. 2025). The resulting accumulation of aberrant proteins disrupts cellular proteostasis, activates protein quality-control pathways, and ultimately compromises growth and development (Thives Santos et al. 2024; Demash et al. 2025; Alles et al. 2026).

In nature, Aze is produced and accumulated by several plant species, in which it is thought to function as a chemical defense (Bell et al. 2008; Gibson et al. 2025). By disrupting protein synthesis in susceptible organisms, Aze inhibits the growth of surrounding organisms, deters herbivores, and suppresses microbial pathogens (Bell et al. 2008; Demash et al. 2025; Rodgers et al. 2025). Its accumulation may therefore enhance plant fitness by reducing competition and limiting damage caused by herbivores and pathogens. Although Aze is toxic to most organisms, several microorganisms that grow in close association with Aze-producers have evolved enzymatic mechanisms for Aze detoxification and assimilation. In *Saccharomyces cerevisiae*, for example, the Aze-specific N-acetyltransferase Mpr1 converts Aze to N-acetyl-Aze, which is no longer recognized by Pro-tRNA synthetase and not misincorporated during protein synthesis (Takagi et al. 2000; Shichiri et al. 2001). The filamentous fungus *Aspergillus nidulans* employs a similar detoxification system comprising the N-acetyltransferase NgnA and the haloacid dehalogenase-like hydrolase AzhA (Biratsi et al. 2021). AzhA mediates hydrolytic opening of the azetidine ring, enabling the subsequent assimilation of Aze through the γ-aminobutyric acid catabolic pathway. Similarly, *Pseudomonas* sp. strain A2C produces an L-azetidine-2-carboxylate hydrolase (HAS) that cleaves the azetidine ring to generate 2-hydroxy-4-aminobutyrate (HAB), initiating a catabolic pathway that allows Aze to serve as a carbon and nitrogen source (Gross et al. 2008). Collectively, these studies demonstrate multiple pathways through which microorganisms have evolved tolerance to Aze. Based on these findings, we investigated whether bacterial HAS could provide Aze tolerance in plants.

To validate our metabolic engineering strategy, we generated transgenic *Arabidopsis thaliana* constitutively expressing a codon-optimized bacterial HAS. We demonstrate that the introduced enzyme localizes to the cytosol and catalyzes the stereoselective degradation of L-Aze *in planta* and *in vitro*. Transgenic plants exhibited markedly enhanced Aze tolerance, maintaining root elongation, cotyledon development, and survival at concentrations that severely inhibited wild-type growth. Integrated transcriptomic and misincorporation proteomic analyses identified the principal molecular pathways underlying Aze toxicity and revealed the cellular mechanisms associated with hydrolase-mediated tolerance. Together, these findings establish hydrolytic Aze degradation as an effective strategy for protecting plants against a toxic amino acid analogue and provide proof of concept that microbial catabolic enzymes can be transferred into plants to expand their metabolic detoxification capacity.

## Results

### Hydrolase expression in Arabidopsis confers Aze tolerance

Previous studies demonstrated that bacterial HAS detoxifies Aze by conversion into the non-toxic metabolite HAB (Gross et al. 2008). We hypothesized we could hijack this bacterial detoxification mechanism to confer Aze tolerance in plants, thus we generated transgenic Arabidopsis plants constitutively expressing the bacterial HAS under the control of the CaMV 35S promoter (Fig. S1). Among the independent transgenic lines examined, the homozygous T3 H-10 line exhibited the greatest Aze tolerance and was therefore selected for subsequent phenotypic analyses (Fig. S2). When exposed to 100 µM Aze, H-10 plants displayed markedly improved overall growth compared with wild-type (WT) plants (Fig. 1A). We next quantified hypocotyl length across increasing Aze concentrations. WT seedlings exhibited pronounced growth inhibition, with reductions of approximately 80% and 90% at 100 and 250 µM Aze, respectively (Fig. 1A, B). In contrast, H-10 seedlings maintained significantly greater growth, with reductions of only approximately 30% and 50% at 100 and 250 µM Aze, respectively (Fig. 1A, B). H-10 plants remained viable at Aze concentrations of up to 250 µM, further demonstrating their enhanced tolerance to Aze (Fig. 1A). Growth on a range of Aze concentrations enabled calculation of IC_50_ values of root inhibition for quantitative comparisons. Similar to previous studies (Thives Santos et al. 2024), WT controls showed an IC_50_ of around 5.22 μM, whereas H-10 showed a significant increase to 6.81 μM Aze (Fig. 1C). We next performed a recovery assay to compare the survivability of WT and H-10 seedlings following exposure to a high concentration of Aze. Seedlings were grown for 5 days on Aze-free medium, transferred to medium containing 500 µM Aze for 8 days, and then returned to Aze-free medium to assess their ability to recover. Nearly all H-10 seedlings resumed growth following removal of Aze, whereas WT seedlings failed to recover, resulting in 0% survival (Fig. 1D and Fig. S3). Thus, HAS expression provides robust protection against acute Aze toxicity and substantially improves seedling recovery.

**Fig. 1.**
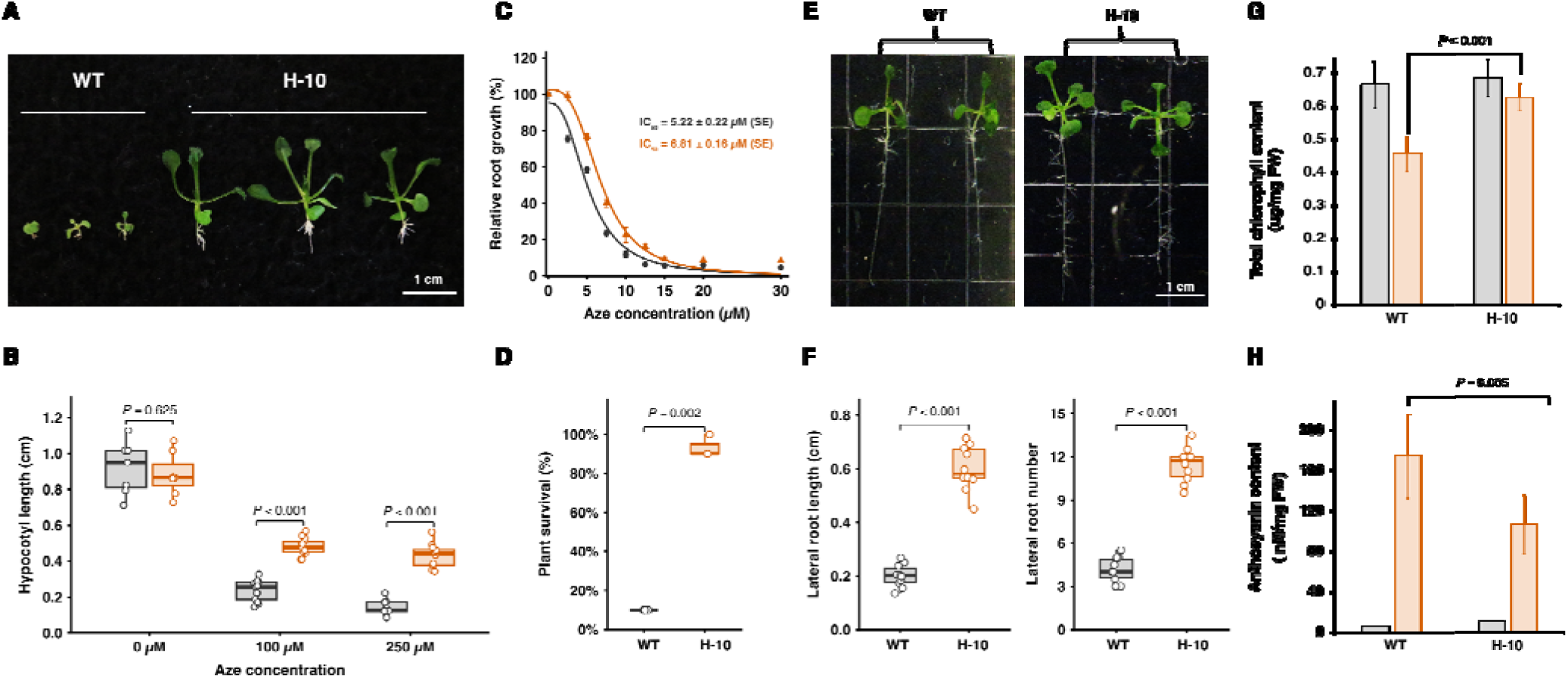
HAS overexpression in Arabidopsis enhances Aze tolerance and mitigates stress responses. (A) Representative images of 14-d-old WT and HAS-overexpressing (H-10) Arabidopsis seedlings grown on medium containing 250 µM azetidine-2-carboxylic acid (Aze). (B) Hypocotyl length of WT (gray) and H-10 (orange) seedlings after 8 d of growth on medium containing 0 µM Aze (*n* > 10), 100 µM Aze (*n* > 12), or 250 µM Aze (*n* > 11). Boxes represent the interquartile range, center lines indicate the median, whiskers extend to 1.5 times th interquartile range, and individual points represent individual seedling measurements. Statistical significance was determined using a two-tailed Welch’s *t*-test; *P* < 0.001. (C) Dose–response analysis of Aze-induced root-growth inhibition in WT (black) and H-10 (orange) seedlings. Relative root growth was measured across increasing Aze concentrations and normalized to the corresponding untreated control. Symbols represent mean ± standard error of the mean, and solid lines represent fitted nonlinear dose–response curves used to estimate the half-maximal inhibitory concentration (IC ). Calculated IC_50_ values are shown. (D) Quantification of WT (gray) and H-10 (orange) plant survival after growth on medium containing 500 µM Aze for 8 d. Individual points represent three independent biological replicates, each comprising 10 plants per genotype. Statistical significance was determined using a two-tailed Welch’s t-test (P = 0.002). (E) Representative images showing the effects of 50 µM Aze on lateral-root development. (F) Quantification of lateral-root length, calculated as the mean length of all lateral roots per seedling and lateral-root number in WT (gray) and H-10 (orange) seedlings grown on medium containing 50 µM Aze (n = 10 seedlings per genotype). Boxes represent the interquartile range, center lines indicate the median, whiskers extend to 1.5 times the interquartile range, and individual points represent individual seedling measurements. Statistical significance was determined using a two-tailed Welch’s *t*-test; *P* < 0.001. (G) Chlorophyll content in WT (gray) and H-10 (orange) under 0 and 100 µM Aze treatments. Bars represent mean of 5 pooled biological replicates ± standard deviation. Statistical significance was determined using a two-tailed Welch’s *t*-test; *P* < 0.001. (H) Anthocyanin content in WT (gray) and H-10 (orange) under 0 and 100 µM Aze treatments. Bars represent mean of 4 pooled biological replicates ± standard deviation. Statistical significance was determined using a two-tailed Welch’s *t*-test; *P* = 0.005.

In Arabidopsis, Aze induces lateral root formation and triggers aboveground stress phenotypes such as reduced chlorophyll content and accumulation of anthocyanins (Alles et al. 2026). We hypothesized that H-10 lines would show a reduced stress phenotype, consistent with increased Aze tolerance. To test this hypothesis, we grew WT and H-10 lines on varying Aze concentrations and assessed lateral root formation, chlorophyll and anthocyanin contents. To determine the effects of Aze on lateral roots, seedlings were grown for 5 days on Aze-free media, then transferred to media containing different concentrations of Aze. Under these conditions, H-10 seedlings developed significantly more lateral roots and maintained greater lateral root growth than WT seedlings (Fig. 1E, F). Aze treatment caused pronounced chlorosis and a substantial reduction in chlorophyll content in WT plants (Fig. 1G). By contrast, H-10 plants retained significantly higher chlorophyll levels and exhibited less visible chlorosis under the same conditions (Fig. 1G). Aze-induced anthocyanin accumulation was also lower in H-10 than in WT plants, consistent with reduced physiological stress (Fig. 1H). Together, these findings indicate that HAS expression alleviates the adverse effects of Aze on plant growth and reduces associated stress responses.

Finally, we investigated whether constitutive expression of HAS imposed a growth penalty in the absence of Aze by growing the plants in soil. H-10 and WT seeds exhibited comparable germination rates under control conditions, indicating that HAS expression did not impair germination or early seedling establishment. During later vegetative development, however, H-10 plants showed slightly slower growth and produced narrower, more elongated leaves than WT plants (Fig. S4). Apart from these modest morphological differences, no major developmental abnormalities were observed. These results demonstrate that heterologous expression of bacterial HAS confers strong Aze tolerance while having only a limited effect on plant growth under non-stress conditions. Collectively, our findings establish enhanced Aze catabolism as an effective metabolic-engineering strategy for protecting plants against the toxic effects of this non-proteogenic amino acid.

### Subcellular localization of HAS

The bacterial HAS lacks an apparent organelle-targeting sequence and was therefore expected to localize to the cytosol. Consistent with this expectation, TargetP and DeepLoc-1.0 identified no N-terminal transit peptide and predicted a cytosolic localization (Fig. S5A). To experimentally validate these predictions, a translational fusion of HAS with yellow fluorescent protein (HAS– YFP) was transiently expressed in *Arabidopsis* protoplasts and *Nicotiana benthamiana* leaf epidermal cells. In both expression systems, the HAS–YFP signal was distributed throughout the cytosol and overlapped with free red fluorescent protein (RFP) signal, which served as a cytosolic marker (Fig. 2A and Fig. S5B). Free YFP was included as an additional control and exhibited a fluorescence pattern comparable to that of HAS–YFP, including overlap with the RFP signal (Fig. 2A). Together, these observations demonstrate that heterologously expressed HAS predominantly localizes to the cytosol in Arabidopsis.

**Fig. 2.**
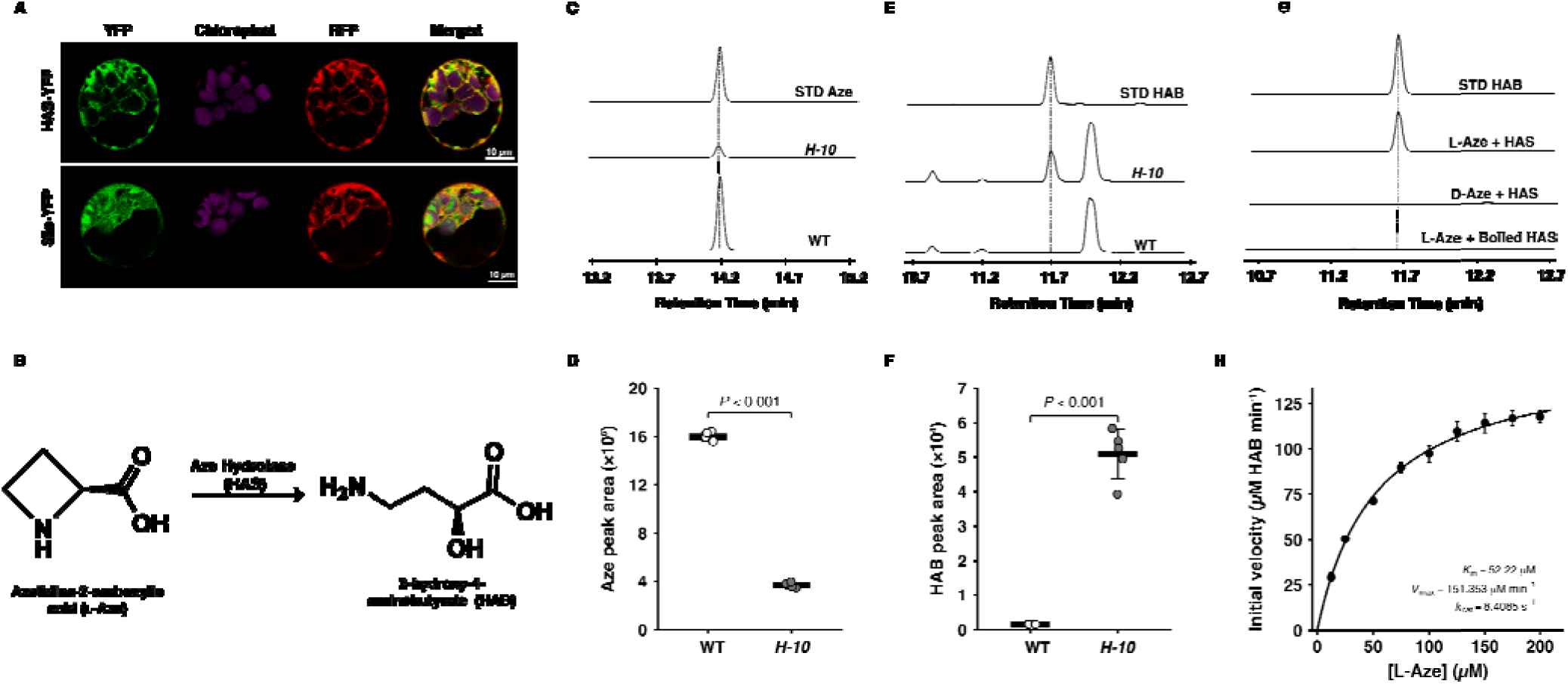
HAS catabolism of Aze to HAB *in planta* and *in vitro.* (A) Representative confocal images of *Arabidopsis thaliana* mesophyll protoplasts transiently expressing HAS-YFP or free YFP. Free RFP was coexpressed as a cytosolic marker. YFP fluorescence is shown in green, chlorophyll autofluorescence in magenta, and RFP fluorescence in red. Colocalization of HAS–YFP with the cytosolic RFP marker is shown in the merged images. Scale bars, 10 µm. Similar cytosolic localization was observed when HAS-YFP wa transiently expressed in *N. benthamiana* (Fig. S5). (B) Schematic representation of the HAS-catalyzed hydrolytic cleavage of L-Aze to produce 2-hydroxy-4-aminobutyrate (HAB). Representative LC–MS/MS chromatograms (C) and quantification (D) of Aze from metabolite extracts of WT and H-10 seedlings treated with 100 µM L-Aze and authentic standards (STD). Individual points represent independent biological replicates ( ); horizontal lines and error bars indicate the mean ± standard deviation. Statistical significance was determined using a two-tailed Welch’s *t*-test; *P* < 0.001. Representative LC–MS/MS chromatograms (E) and quantification (F) of HAB from metabolite extracts of WT and H-10 seedlings treated with 100 µM L-Aze and authentic standards (STD). Individual points represent independent biological replicates ( ); horizontal lines and error bars indicate the mean ± standard deviation. Statistical significance was determined using a two-tailed Welch’s *t*-test; *P* < 0.001. (G) Representative LC–MS/MS chromatograms showing HAB formation in reactions containing purified HAS and L-Aze. HAB was not detected in reactions containing D-Aze or heat-inactivated HAS. (H) Michaelis–Menten kinetics of purified HAS using L-Aze as the substrate. Points represent mean initial velocities from two independent replicates, error bars indicat standard error of the means, and the solid line represents the nonlinear Michaelis–Menten fit. The estimated kinetic parameters were *k_m =_* 52.22 *μ*M and *k*cat = 8.4085 s^−1^ (*R*^2^ = 0.99).

### HAS catalyzes the conversion of Aze to HAB *in vitro* and *in planta*

We next investigated whether the enhanced Aze tolerance of H-10 plants was associated with the enzymatic conversion of Aze to HAB *in planta*. Fig. 2B schematically illustrates the HAS-catalyzed hydrolytic cleavage of L-Aze to produce 2-hydroxy-4-aminobutyrate (HAB). To quantify Aze and HAB accumulation, metabolites were extracted from leaf tissues of WT and H-10 plants treated with 100 µM Aze and analyzed by liquid chromatography–tandem mass spectrometry (LC–MS/MS) and compared with authentic standards. Chromatographic analysis revealed lower Aze abundance in H-10 seedlings than in WT, whereas HAB was detected in H-10 plants and not in WT (Fig. 2C-F). Quantitative analysis confirmed that H-10 plants accumulated substantially less Aze and significantly more HAB than WT plants (Fig. 2D, F). These findings demonstrate that H-10 plants actively convert Aze to HAB *in planta* (Fig. 2C-F) and support the conclusion that HAS-mediated Aze degradation underlies the enhanced Aze tolerance of H-10 plants.

Given the ability of HAS to convert Aze into HAB *in planta*, we wanted to determine its kinetic parameters *in vitro*. For biochemical characterization, recombinant HAS was expressed in *Escherichia coli* and purified by affinity chromatography (Fig. S6A). The purified enzyme converted Aze to HAB, confirming its proposed role in Aze catabolism (Fig. 2G and Fig. S6). To determine the stereospecificity of HAS, we tested both L- and D-Aze as substrates. HAS converted L-Aze to HAB, whereas no activity with D-Aze was detected at concentrations up to 200 µM under identical assay conditions, indicating stereospecificity for L-Aze (Fig. 2G). Heat-denatured HAS failed to produce detectable HAB, confirming that product formation required an active enzyme. Furthermore, HAS displayed no detectable activity toward Pro, indicating that the enzyme evolved strict specificity for L-Aze (Fig. S6D). We next examined the effects of metal ions on HAS activity. None of the ions tested, including K^⁺^, Fe^²⁺^, and Mg²^⁺^, significantly enhanced activity. A representative chromatogram from the Mg²^⁺^ assay is shown in Fig. S6D. Michaelis–Menten kinetic analysis yielded an apparent *K_m_* of 52.22 µM and a *k_cat_* of 8.4 s□¹, corresponding to a catalytic efficiency ( *k_cat_*/*K_m_*) of approximately 1.61 × 10^5^ M□¹ s□¹ (Fig. 2H).

### Hydrolase expression attenuates Aze-induced proteotoxic stress and preserves growth-associated pathways

To investigate the molecular mechanisms underlying the enhanced Aze tolerance of H-10, we performed RNA sequencing of WT and H-10 seedlings grown in the absence or presence of 100 μM Aze. Principal component analysis (PCA) clearly separated the treatment groups, with PC1 accounting for 93.2% of the total variance (Fig. S7A). Aze-treated WT samples were strongly separated from all other groups, indicating extensive Aze-induced transcriptomic reprogramming (Fig. S7A). In contrast, untreated WT and H-10 samples clustered closely (Fig. S7A). Aze-treated H-10 samples remained more similar to the untreated groups (Fig. S7A). Pearson correlation analysis confirmed high reproducibility among biological replicates and a strong similarity between untreated WT and H-10 samples (Fig. S7B). Moreover, Aze-treated H-10 samples retained higher correlations with untreated samples than did Aze-treated WT samples (Fig. S7B).

Next, to characterize the differential transcriptional responses of WT and H-10 to Aze, we compared gene expression between Aze-treated H-10 and WT seedlings. Hierarchical clustering of the top 100 differentially expressed genes (DEGs) revealed a prominent cluster that was strongly expressed in WT but showed substantially lower expression in H-10 (Fig. 3A). Conversely, a smaller cluster of genes was maintained at higher levels in H-10 than in WT. Functional enrichment analysis showed that genes preferentially expressed in WT were associated with cellular stress, oxidative stress, defense responses, and the unfolded protein response (UPR) (Fig. 3B, C). In contrast, genes expressed at higher levels in H-10 were enriched in photosynthesis, carbon fixation, plastid function, cellular growth, and energy metabolism (Fig. 3B, C). These patterns indicate that WT seedlings activate a broad stress-response program following Aze exposure, whereas H-10 maintains transcriptional programs associated with photosynthetic activity, energy production, and growth. Because our previous studies demonstrated that Aze misincorporation induces protein misfolding and activates the unfolded protein response (UPR) (Thives Santos et al. 2024; Alles et al. 2026), we specifically examined genes associated with the UPR and endoplasmic reticulum protein-quality control. Key UPR components, including *BIP1*, *BIP2, BIP3*, *ERDJ2A*, *ERDJ3A*, *CRT1A*, *CNX1*, and *IRE1-2*, were strongly induced in Aze-treated WT seedlings but showed substantially weaker induction in H-10 (Fig. 3D). This attenuated UPR signature in response to Aze is consistent with reduced proteotoxic stress in H-10 (Fig. 4).

**Fig. 3.**
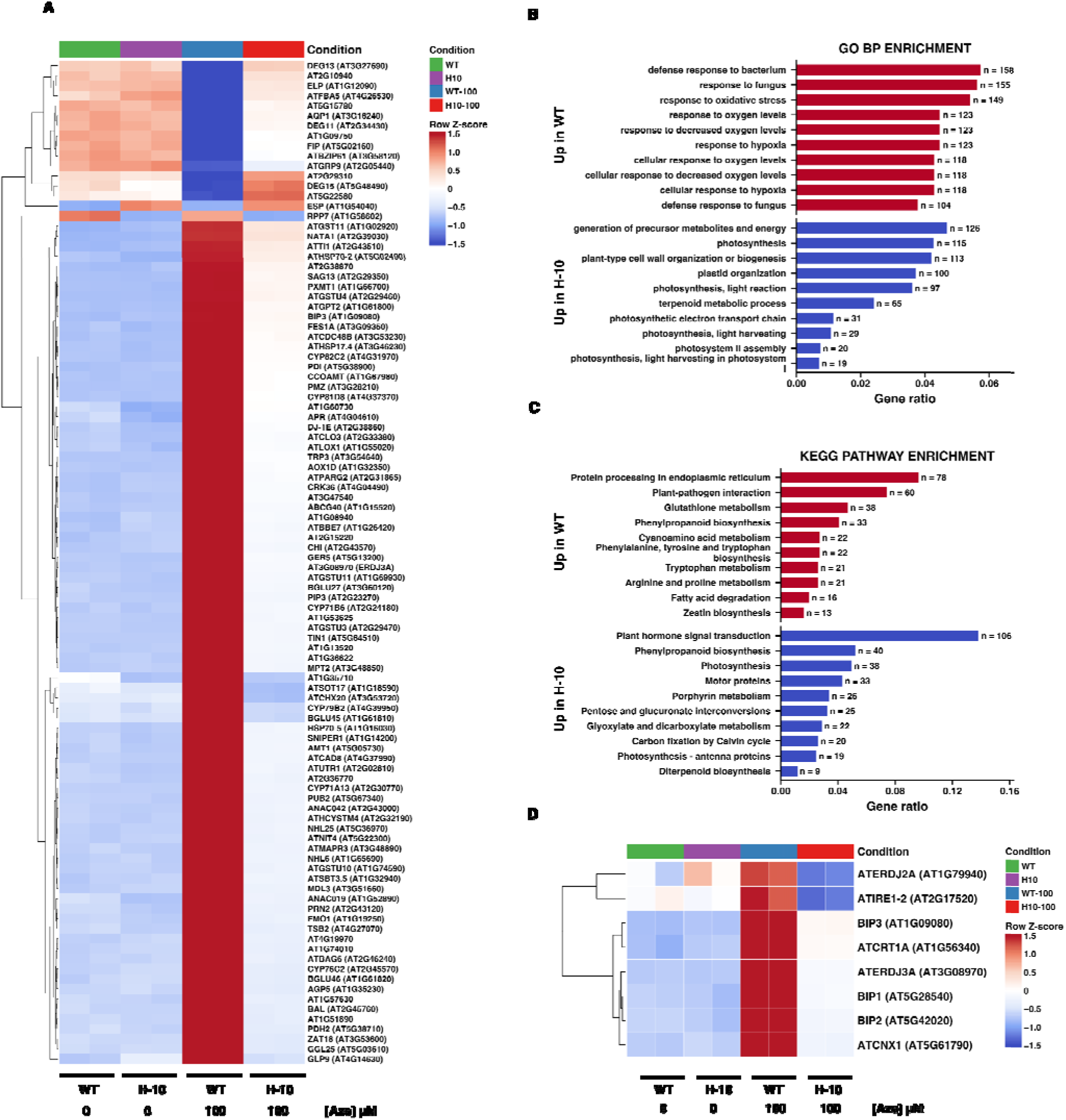
H-10 represses Aze-induced stress responses and preserves photosynthetic functions at the transcriptional level. RNA-seq analysis of WT and H-10 seedlings grown for 8 d on medium containing 0 or 100 µM Aze. (A) Hierarchically clustered heatmap of the top 100 differentially expressed genes (DEGs) identified between WT and H-10 seedlings treated with 100 µM Aze. Expression patterns are shown across untreated and Aze-treated WT and H-10 seedlings, with two independent biological replicates per condition. Colors represent row-wise Z-scores calculated from variance-stabilized expression values, with red and blue indicating higher and lower relative expression, respectively. (B, C) Functional enrichment analyses of DEGs between WT and H-10 seedlings treated with 100 µM Aze. (B) Gene Ontology Biological Process (GO BP) enrichment analysis. (C) Kyoto Encyclopedia of Genes and Genomes (KEGG) pathway enrichment analysis. Red bars indicate terms enriched among genes expressed at higher levels in WT treated with Aze, whereas blue bars indicate terms enriched among genes expressed at higher levels in H-10 treated with Aze. Gene ratio represents the proportion of DEGs in the corresponding directional gene list assigned to each term or pathway, and values adjacent to the bars indicate the number of assigned genes (n). GO terms and KEGG pathways with adjusted *P* < 0.05 were considered significantly enriched. (D) Hierarchically clustered heatmap showing the expression of selected genes associated with the unfolded protein response and endoplasmic reticulum protein quality control. Colors represent row-wise Z-scores calculated from variance-stabilized expression values. These genes were strongly induced in WT following Aze treatment, whereas their induction was attenuated in H-10.

**Fig. 4.**
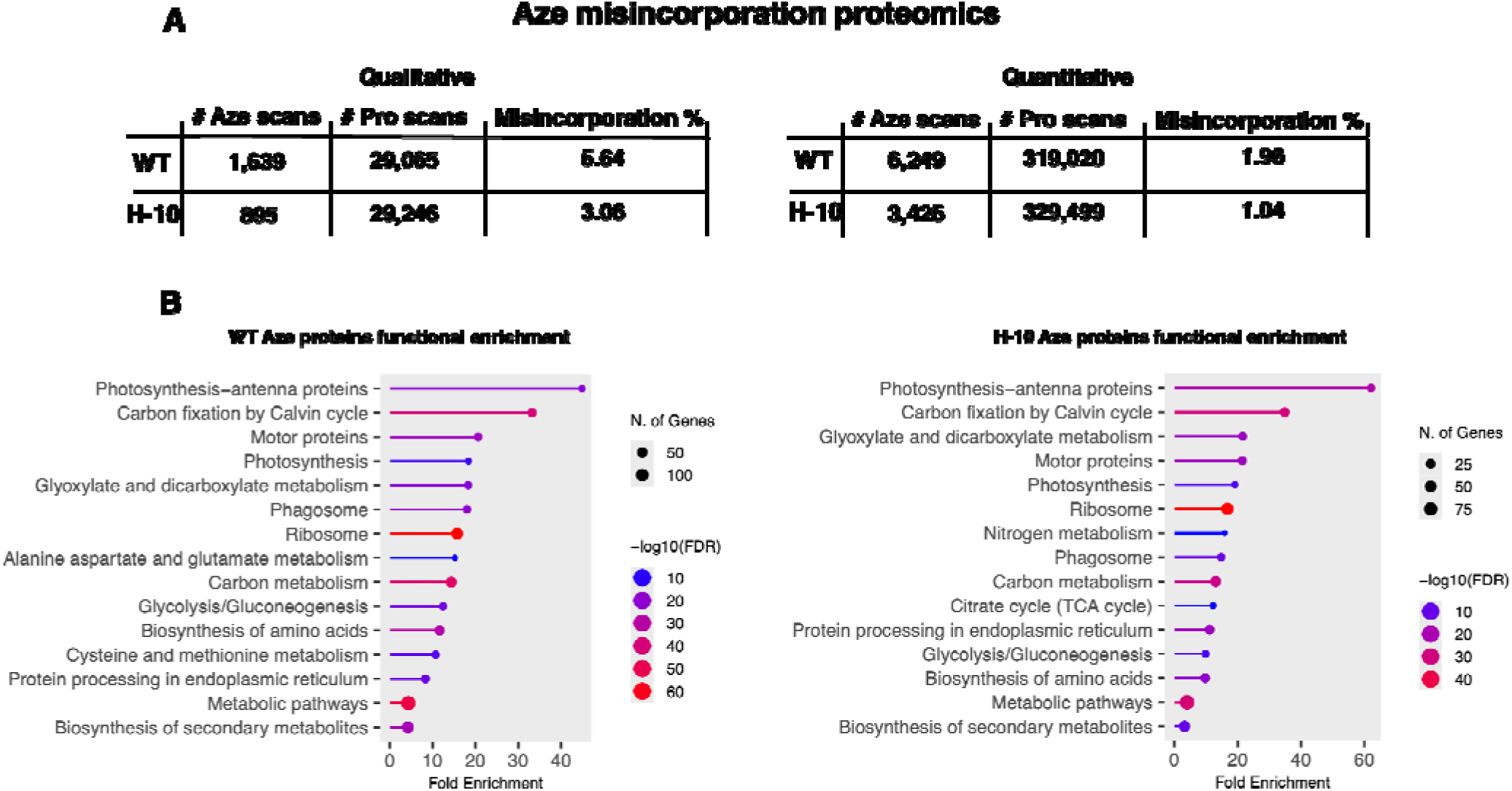
Proteomic analysis reveals reduced Aze misincorporation in H-10. (A) Qualitative and quantitative comparison of Aze misincorporation events detected across the proteomes of WT and H-10 seedlings following 100 μM Aze treatment for 8 days. Aze misincorporation events were identified by treating Aze as a post-translational modification of Pro. Qualitative data represents the number of peptides with Aze and the number of peptide detected with a Pro. Whereas quantitative reflects the number of Aze or Pro containing peptide and the number of scans for each of those peptide (B) Kyoto Encyclopedia of Genes and Genomes (KEGG) functional enrichment analysis of proteins with Aze misincorporation in Aze-treated WT and H-10 seedlings. Similar pathways and processes were functionally enriched. However, misincorporation rates were reduced in H-10.

To assess pathway-level changes independently of a predefined DEG threshold, we performed gene-set enrichment analysis (GSEA) using the complete gene list ranked by differential expression between H-10 and WT plants treated with 100 µM Aze (H-10-100 versus WT-100; Fig. S8). Positive normalized enrichment scores (NESs) indicated enrichment in 100 µM Aze H-10, whereas negative NESs indicated enrichment in 100 µM Aze WT. GO Biological Process GSEA revealed enrichment of photosynthetic and chloroplast-organization processes in H-10, while protein catabolism, vacuolar transport, fungal responses, and ER stress were enriched in WT (Fig. S8A). KEGG GSEA similarly showed enrichment of photosynthesis, porphyrin metabolism, ribosome, carbon fixation, and carbon metabolism in H-10, whereas ER protein processing, proteasome, glutathione metabolism, plant–pathogen interaction, spliceosome, and endocytosis were enriched in WT (Fig. S8B). The agreement between GSEA and DEG-based enrichment analyses supports the conclusion that H-10 maintains photosynthetic and metabolic functions during Aze exposure, whereas WT undergoes pronounced proteotoxic and cellular stress. Consistent with the greater anthocyanin accumulation observed in Aze-treated WT plants (Fig. 1H), we next examined the expression of key genes involved in phenylpropanoid and flavonoid biosynthesis (Fig. S9). *PAL1*, *PAL2*, *C4H*, *4CL2*, *CCR1*, *HCT*, *CHS*, *ANS*, *CYP75B1*, and *UGT78D2* were strongly induced in WT but showed weaker or no induction in H-10. Analysis of the 48 most variable ER stress-associated genes further revealed coordinated activation of the ER protein-quality-control machinery in WT (Fig. S10). This response included genes encoding UPR regulators (*bZIP17*, *bZIP28*, *bZIP60*, *ANAC062*, and *ANAC089*), ER chaperones and co-chaperones (*SDF2*, *P58IPK*, *ERDJ3B*, and *HSP90-7*), as well as genes involved in glycoprotein quality control, protein folding and redox regulation, ER-associated degradation, protein translocation, and cell-fate regulation. In contrast, most of these genes exhibited markedly weaker induction in H-10, indicating that HAS expression attenuates Aze-induced activation of ER stress and protein-quality-control pathways. Collectively, the diverse gene expression analyses demonstrate that WT seedlings undergo extensive transcriptomic reprogramming following Aze exposure upregulating proteotoxic stress pathways, while H-10 maintains energy metabolism.

### Proteomic analysis reveals reduced Aze misincorporation in H-10

We hypothesized that enhanced Aze tolerance of H-10 results from HAS-mediated depletion of intracellular Aze limiting its misincorporation into newly synthesized proteins. We performed misincorporation proteomics for WT and H-10 grown on 100 μM Aze for 8 days to determine if enhanced Aze tolerance in H-10 is due to reduced misincorporation rates. In WT, a total of 29,065 peptides were identified with ≥2 scans across the three biological replicates at a 1% false discovery rate (Fig. 4A). A similar number of peptides were identified in H-10, 29,246 (Fig. 4A). Next, Aze misincorporation events were identified by treating Aze as a post-translational modification with a mass difference of 14.01 *m/*z between Pro and Aze. Peptides with a Pro to Aze modification with an Ascore ≥20 were treated as misincorporation events. The data were then analyzed in two ways, one which took into the account the peptide scan numbers, and another that was qualitative and treated Aze misincorporation as a yes/no. In WT, 1,639 peptides were identified with an Aze misincorporation rate of 5.64% (Fig. 4A). The misincorporation rate for H-10 of 3.06% was reduced by 45% compared to WT (Fig. 4A). These misincorporation rates are consistent with a previous misincorporation study of Arabidopsis grown on Aze (Thives Santos et al., 2024). Considering peptide scan numbers, Aze misincorporation rates were lower, however a consistent reduction of Aze misincorporation rate in H-10 was observed by around 47% (Fig. 4A). These data suggest that the increased Aze tolerance of H-10 is due to its substantially reduced Aze misincorporation rates.

Not only could the proteomics data be used to quantify misincorporation rates, but also the proteomic responses to Aze in WT and H-10. In WT, we find functional enrichment of categories associated with the phagosome, ribosome, and protein processing at the ER (Fig. 4B). These processes are consistent with the transcriptomic analysis and a previous study that investigated the proteomic response to Arabidopsis grown on Aze (Fig. 3 and Figs. S8 and S10), (Thives Santos et al. 2024). Similar to the transcriptomic response, core metabolic processes and photosynthesis and related terms are enriched in the proteomics data (Fig. 4B), again consistent with the ability of H-10 to maintain energy metabolism instead of triggering broad proteotoxic stress responses. Functional enrichment analysis of proteins with Aze misincorporation in H-10 showed enrichment of similar processes compared to WT including the phagosome, ribosome, and protein processing at the ER (Fig. 4B). We next looked at proteins that were more abundant in WT compared to H-10. Functional enrichment analysis was performed using proteins with a 2-fold increased abundance in WT (consisting of 321 proteins). Enriched processes related to ATP hydrolysis, RNA binding and hydrolase activity (Fig. S11A). These processes indicate stress and abnormalities in transcription and translation (Fig. S11A). We next took a targeted analysis of proteins involved in the unfolded protein response, as this process is upregulated in response to Aze treatment. Most of the proteins involved in the UPR were more abundant in WT than in H-10 or showed little change (Fig. S11B) suggesting a diminished proteotoxic stress response due to lower misincorporation rates, consistent with the transcriptomic response.

## Discussion

Aze is produced by several plant species as a defense metabolite that deters herbivores, restricts microbial growth, and suppresses growth of surrounding organisms (Thives Santos et al. 2024; Demash et al. 2025; Gibson et al. 2025; Rodgers et al. 2025). The ability of Aze to disrupt proteostasis in both plant and mammalian systems underscores the importance of elucidating its metabolism and detoxification, both for evaluating potential dietary risks and for developing strategies to limit its accumulation in food crops (Song et al. 2017; Thives Santos et al. 2024; Demash et al. 2025; Rodgers et al. 2025). To determine whether Aze could be enzymatically detoxified *in planta*, we expressed a bacterial hydrolase belonging to the haloacid dehalogenase superfamily in Arabidopsis. This enzyme hydrolytically opens the azetidine ring of Aze to generate HAB (Fig. 2B), converting a toxic Pro analog into a non-toxic product that does not compete with Pro during protein biosynthesis. Hydrolase-expressing Arabidopsis exhibited substantially greater tolerance and survivability to Aze than WT plants (Fig. 1). Although the ecological function of Aze remains unknown, the emergence of Aze-degrading or -modifying enzymes in bacteria and yeast reiterates its ecological relevance.

Subcellular localization provides a mechanistic explanation for the protective activity of bacterial HAS. Arabidopsis contains two Pro-tRNA synthetases (ProRS), the enzymes responsible for recognizing and activating Pro during protein biosynthesis (Duchêne et al. 2005; Lee et al. 2016). The cytosolic Pro-tRNA synthetase recognizes Aze and Pro equally (Lee et al. 2016) and thus is responsible for misincorporation of Aze for Pro during translation, whereas the organellar localized Pro-tRNA synthetase prefers Pro to Aze by about 100-fold (Lee et al. 2016). The bacterial hydrolase localized predominantly to the cytosol, which is a principal site of protein biosynthesis (Fig. 2A). This localization places the enzyme in an appropriate cellular compartment to degrade Aze before it is activated by cytosolic Pro-tRNA synthetase. LC– MS/MS analysis revealed lower Aze abundance and substantially greater HAB accumulation in H-10 plants than in WT plants following Aze treatment (Figs. 2C-F). Furthermore, proteomic analysis revealed an approximately 50% reduction in Aze misincorporation rates in H-10 compared with WT (Fig. 4). These results support a model in which cytosolic Aze degradation decreases the intracellular Aze pool available for aminoacylation, particularly in the cytosol, thereby limiting its misincorporation into newly synthesized proteins.

Biochemical characterization of recombinant HAS established the catalytic basis of the enhanced tolerance observed in H-10 plants. Purified HAS converted Aze into HAB (Figs. 2G, H and Fig. S6). Kinetic analysis yielded a *K*_m_ of 52.22 µM and a *k_cat_* of 8.4 s ¹ indicating effective substrate recognition and catalytic turnover within the concentration range relevant to the plant-tolerance assays (Fig. 2G, H and Fig. S6). The enzyme also exhibited stereoselectivity and substrate specificity: L-Aze was converted to HAB, whereas no detectable activity was observed with D-Aze or L-Pro (Fig. 2G). It is interesting that HAS is capable of distinguishing Aze from Pro, whereas ProRS, which are ancient enzymes used in translation are unable to discriminate between Aze and Pro in the primary active site (Chaliotis et al. 2017), whereas HAS is a more recent evolutionary innovation, it has quickly evolved strict specificity. Although, if HAS was not strictly specific towards Aze, then it would catabolize Pro leading to less Pro available for protein biosynthesis. Understanding how HAS distinguishes Aze from Pro at the atomic level may provide insights into broader classes of enzymes.

The transcriptomic and proteomic results further support a detoxification-based mechanism. Transcriptomic analyses provide a mechanistic framework for the enhanced Aze tolerance of H-10 (Fig. 3). Aze caused extensive transcriptional reprogramming in WT, characterized by strong activation of the UPR, ER protein processing, proteasomal degradation, antioxidant defenses, and other stress pathways (Fig. 3 and Figs. S7 and S8). These responses are consistent with Aze misincorporation disrupting protein folding and imposing substantial proteotoxic and oxidative stress (Thives Santos et al. 2024; Alles et al. 2026). The coordinated induction of phenylpropanoid and anthocyanin biosynthetic genes further reflects the broad stress response of WT seedlings (Fig. 1 and Fig. S9). Notably, toxicity caused by another non-proteogenic amino acid, meta-tyrosine, has similarly been associated with perturbations in phenylpropanoid metabolism (Zer et al. 2020), whereas canavanine disrupts reactive oxygen and nitrogen species homeostasis (Staszek and Gniazdowska 2020). These parallels raise the possibility that structurally distinct non-proteogenic amino acids activate overlapping stress-response pathways following misincorporation. By contrast, these stress-associated transcriptional responses were markedly attenuated in H-10, further supporting the conclusion that HAS expression reduces Aze-induced cellular stress while preserving core metabolic processes (Fig. 3 and Figs. S7 and S8). Proteomic analysis provided more direct evidence that HAS limits the entry of Aze into protein biosynthesis. Under Aze stress, H-10 plants exhibited an approximately 50% reduction in Aze misincorporation relative to WT plants (Fig. 4), consistent with enzymatic depletion of the cytosolic Aze pool and reduced competition between Aze and Pro for recognition and aminoacylation. The persistence of some Aze misincorporation in H-10 may reflect competition between Aze hydrolysis and aminoacylation. Kinetic analysis indicated that cytosolic ProRS has a lower apparent substrate affinity for Aze than HAS (40 µM versus 52.22 µM for HAS; Lee et al., 2016). Because both enzymes are cytosolic, competition for the available Aze may explain the residual Aze misincorporation observed in H-10. Although HAS expression provides a promising strategy for Aze detoxification in plants, the molecular basis for its recognition of L-Aze over Pro and D-Aze remains unclear. Analysis of the L-Aze-soaked HAS crystal structure (PDB 8YWO) revealed a well-defined active-site pocket formed by Asp12, Tyr14, Trp20, Tyr64, and Asn122 (Fig. S12). Asp12 is positioned near the cleaved C–N bond. Consistent with a role in substrate positioning or catalysis; notably, the D12A substitution was previously shown to abolish enzymatic activity (Gross et al. 2008). The remaining pocket residues may stabilize L-Aze in a catalytically productive orientation while disfavoring Pro and D-Aze. However, these proposed interactions and the structural basis of stereoselectivity require validation by site-directed mutagenesis and biochemical analysis. Defining the relative kinetics and intracellular competition between HAS-mediated hydrolysis and ProRS-mediated activation will clarify how effectively HAS intercepts Aze before its incorporation into protein biosynthesis. These mechanistic insights could ultimately guide targeted mutagenesis or directed evolution of HAS to enhance its catalytic efficiency and further improve Aze tolerance in plants.

Collectively, our findings establish HAS-mediated hydrolysis as an effective strategy for Aze detoxification that preserves cellular proteostasis and sustains plant growth under Aze stress. This approach may provide a framework for protecting crops from Aze-mediated proteotoxicity and could be extended to other toxic non-proteogenic amino acids when suitable substrate-specific enzymes are available. Future engineering strategies could integrate spatially or developmentally regulated Aze biosynthesis with tissue-specific detoxification, thereby balancing the defensive benefits of this specialized metabolite with the protection of essential cellular processes.

## Materials and Methods

### Generation of transgenic *Arabidopsis thaliana* expressing the Aze hydrolase

The coding sequence of the bacterial azetidine-2-carboxylic acid (Aze) hydrolase (HAS) was codon-optimized, commercially synthesized, placed under the control of the cauliflower mosaic virus 35S (CaMV 35S) promoter, and cloned into a binary vector (Fig. S13). The resulting construct was verified by Sanger sequencing and introduced into *Agrobacterium tumefaciens* strain GV3101. Wild-type (WT) *Arabidopsis thaliana* accession Columbia-0 (Col-0) plants were transformed using the floral-dip method (Clough and Bent 1998). Primary transformants were selected on half-strength Murashige and Skoog (½ MS) medium supplemented with 15 mg L ¹ hygromycin. Putative transformants were screened by PCR using primers specific to the hydrolase transgene and the hygromycin resistance gene (Table S1). Independent homozygous T3 lines were established, and line H-10, which exhibited stable hydrolase expression and enhanced Aze tolerance, was selected for subsequent physiological, biochemical, and molecular analyses.

### Plant growth conditions and seed sterilization

Seeds of WT (Col-0) and transgenic line H-10 were surface sterilized by exposure to chlorine gas generated from 150 mL commercial bleach and 4.5 mL 12 M hydrochloric acid in a sealed desiccator for 1 h (Lindsey Iii et al. 2017). Sterilized seeds were sown on ½ MS medium (PhytoTech Labs, USA) containing 1% sucrose and 0.8 % agar concentration. Plates were stratified at 4 °C in darkness for 48 h and then transferred to controlled growth conditions maintained under a 16-h-light/8-h-dark photoperiod at 22 °C and a photosynthetic photon flux density of approximately 150 μmol m□² s□¹.

### Seedling growth assays under Aze stress

To evaluate Aze tolerance, WT and H-10 seeds were sown on ½ MS medium containing 0, 100, or 250 µM L-azetidine-2-carboxylic acid (L-Aze; Sigma-Aldrich). Plates were positioned vertically under controlled growth conditions. After 8 d, seedlings were photographed, and primary-root and hypocotyl lengths were measured using ImageJ. Growth measurements were used to compare the sensitivity of WT and H-10 seedlings to L-Aze. For lateral-root analysis, WT and H-10 seedlings were initially grown vertically on Aze-free ½ MS medium for 5 d. Seedlings of uniform size were transferred to fresh ½ MS medium containing 0, 50, or 100 µM L-Aze. After an additional 5 d of growth, seedlings were imaged, and the number and total length of lateral roots were quantified using ImageJ.

### Aze recovery assay

For recovery assays, WT and H-10 seedlings were initially grown vertically on Aze-free ½ MS medium for 5 d. Seedlings of uniform size were transferred to ½ MS medium containing 500 µM L-Aze and maintained for an additional 5 d. Seedlings were then transferred to Aze-free ½ MS medium and allowed to recover under standard growth conditions for 8 d. Seedling survival was evaluated based on two criteria: the absence of chlorotic leaf tissue and the resumption of growth, as indicated by increased root length. Recovery assays were performed in three independent experiments, each comprising at least three biological replicates.

### Determination of the Aze half-maximal inhibitory concentration

The half-maximal inhibitory concentration (IC□□) of Aze was determined from primary-root growth. WT and H-10 seedlings were grown on ½ MS medium containing 0, 2.5, 5, 7.5, 10, 12.5, 15, 20, and 30 µM L-Aze under the conditions described above. Primary-root length was measured after 8 d and expressed relative to the mean root length of the corresponding genotype at 0 µM Aze. Concentration-response curves were fitted using a four-parameter logistic model, and IC values and their 95% confidence intervals were estimated as described by Sebaugh 2011. At least eight biologically independent seedlings were analyzed at each concentration.

### Chlorophyll and anthocyanin quantification

WT and H-10 seedlings were grown for 8 d on ½ MS medium containing 0 or 100 µM Aze. For chlorophyll extraction, three seedlings were transferred to a 1.5 mL microcentrifuge tube containing 1 mL of 80% acetone and gently rocked at room temperature for 24 h in the dark. Samples were then centrifuged at 15,000 *× g* for 7 min, and the resulting supernatant was used for spectrophotometric analysis. Absorbance was measured at 646 nm and 663 nm and chlorophyll content was measured using equations found in Lichtenthaler 1987. Chlorophyll content was corrected for tissue fresh weight and is the average of 5 pooled biological replicates. For anthocyanin extraction, three seedlings per sample were transferred to a microcentrifuge tube and homogenized thoroughly. The homogenate was resuspended in 400 µL of MeOH/CHCl (2:1 v/v), followed by the addition of 300 µL of H O and 125 µL of CHCl . Samples were vortexed and centrifuged at 15,000 *× g* for 10 min at room temperature to facilitate phase separation. The upper aqueous/methanolic phase, containing the anthocyanins, was carefully collected. To acidify and stabilize the anthocyanin-containing fraction, an equal volume of 0.1 M HCl (300 µL) was added to 300 µL of the collected phase. Absorbance of the resulting solution was measured spectrophotometrically at 520 nm similar to Schenck et al. 2020. Anthocyanin content was corrected for tissue fresh weight and is the average of 4 pooled biological replicates.

### Subcellular localization of Aze hydrolase

To determine the subcellular localization of Aze hydrolase, its coding sequence was fused in frame to yellow fluorescent protein (YFP), generating a C-terminally tagged HAS–YFP fusion under the control of the cauliflower mosaic virus 35S (CaMV 35S) promoter (Table S1). HAS–YFP and free YFP were transiently expressed in *Arabidopsis thaliana* mesophyll protoplasts and *Nicotiana benthamiana* leaves using previously described protocols (Dwivedi et al. 2022, 2026). Free red fluorescent protein (RFP) was coexpressed as a cytosolic marker. Fluorescence was examined using a Leica TCS SP8 confocal laser-scanning microscope 16 h after protoplast transfection or 72 h after leaf infiltration. YFP was excited at 514 nm, and emission was detected at 525–575 nm. RFP was excited at 558 nm, and emission was detected at 570–620 nm. Chlorophyll autofluorescence was recorded separately using excitation at 633 nm and an emission window of 650–725 nm. Images were processed using LAS X software. The subcellular localization of HAS was assessed by comparing the distribution of the HAS–YFP signal with that of free YFP and the cytosolic RFP marker.

### Heterologous expression and purification of the Aze hydrolase

The HAS coding sequence was cloned into the bacterial expression vector pET28b-SUMO to produce an N-terminally SUMO-His-tagged recombinant protein (Table S1). The resulting construct was introduced into *E. coli*. Cultures were grown in LB medium at 37 °C to an OD of approximately 0.6, and protein expression was induced with 0.4 mM isopropyl β-D-1-thiogalactopyranoside (IPTG) at 16 °C for 16 h. Cells were harvested by centrifugation and resuspended in lysis buffer containing 50 mM Tris–HCl (pH 8.0), 500 mM NaCl, 20 mM imidazole, 5% (v/v) glycerol, and 0.1 mg mL ¹ lysozyme. Following disruption by sonication, the lysate was clarified by centrifugation at 10,000 × *g* for 20 min at 4 °C.

The recombinant hydrolase was purified from the soluble fraction by nickel–nitrilotriacetic acid (Ni–NTA) affinity chromatography. The column was washed with buffer containing 50 mM Tris–HCl (pH 8.0), 500 mM NaCl, 20 mM imidazole, and 5% (v/v) glycerol. The bound protein was then eluted with buffer containing 50 mM Tris–HCl (pH 8.0), 500 mM NaCl, 500 mM imidazole, and 5% (v/v) glycerol. The eluate was subsequently desalted and exchanged into buffer containing 100 mM Tris, 150 mM NaCl, and 5% (v/v) glycerol using a centrifugal filter device with a 10-kDa molecular-weight cutoff. Protein concentration was determined using the Bradford assay, and purity was estimated by densitometric analysis of Coomassie-stained SDS– polyacrylamide gels using ImageJ (Fig. S6A).

### Aze hydrolase activity and kinetic analysis

Hydrolase activity was determined by quantifying the formation of 2-hydroxy-4-aminobutyrate (HAB) from L-azetidine-2-carboxylic acid (L-Aze). Standard reactions were performed in a total volume of 120 µL containing 50 mM Tris (pH 8.0), 150 mM NaCl, 0.25 µM purified enzyme, and L-Aze at final concentrations of 0, 12.5, 25, 50, 75, 100, 200, or 400 µM. Reactions were incubated at 30 °C for 5 min and terminated by the addition of acetonitrile. The resulting samples were derivatized with dansyl chloride as described previously (Du et al. 2025). Control reactions lacking enzyme or substrate, as well as reactions containing heat-inactivated enzyme, were processed in parallel.

HAB formation was quantified by liquid chromatography–mass spectrometry (LC–MS) using an authentic HAB standard (Fig. S6B) and an external calibration curve. Product concentrations were corrected using the corresponding zero-time or no-enzyme controls. Initial velocities were calculated by dividing the amount of HAB formed by the reaction time within the experimentally established linear range. Kinetic parameters were estimated by nonlinear regression of initial velocity against L-Aze concentration using the Michaelis–Menten equation. Parameter estimates were reported with their standard errors and 95% confidence intervals.

### Substrate-specificity assay

Substrate specificity was evaluated under standard hydrolase assay conditions using L-Aze, D-Aze, or L-Pro, each at a final concentration of 50 and 200 µM. Product formation was quantified by LC–MS using the corresponding authentic standard where applicable. Activity toward L-Aze was defined as 100%, and activity toward the other substrates was expressed relative to this value. No-enzyme and zero-time controls were included for each substrate and used for background correction.

### Extraction and LC–MS quantification of Aze and HAB from Arabidopsis seedlings

WT and H-10 seedlings were grown for 8 d on half-strength Murashige and Skoog (½ MS) medium containing 0 or 100 µM L-Aze. Whole seedlings were harvested, lyophilized for 16–20 h, and stored at −80 °C until extraction. Each biological replicate consisted of 10–20 pooled seedlings, and five independent biological replicates were analyzed per genotype and treatment. Lyophilized tissue was pulverized with 3-mm glass beads using a bead mill grinder. For metabolite extraction, 2 mg of pulverized dry tissue was mixed with 800 µL of chloroform and incubated at 50 °C for 1 h. Subsequently, 800 µL of water was added, and the samples were incubated again at 50 °C for 1 h. Samples were centrifuged at 3,000 ×g for 20 min at 4 °C to separate the aqueous and organic phases. The upper aqueous phase was carefully transferred to a clean glass vial and lyophilized overnight. The dried extract was reconstituted in 120 µL of water:acetonitrile (1:1, v/v). For derivatization, each reconstituted extract was combined with 10 µL of 1 M sodium borate buffer (pH 8.0) and 10 µL of 20 mM dansyl chloride prepared in acetonitrile (Du et al. 2025). Samples were incubated at 30 °C for 1 h in darkness and subsequently centrifuged at 3,000 ×g for 5 min at room temperature. The resulting supernatants were transferred to autosampler vials for LC–MS analysis. Chromatographic separation was performed using an ACQUITY Premier ultrahigh-performance liquid chromatography system coupled to a Xevo MRT multi-reflecting time-of-flight mass spectrometer (Waters). Dansylated metabolites were separated on an ACQUITY Premier BEH C18 column (2.1 × 100 mm, 1.7-µm particle size, 130-Å pore size) maintained at 40 °C. The mobile phases consisted of water containing 0.1% formic acid (solvent A) and acetonitrile (solvent B). The flow rate was 0.3 mL min ¹, the autosampler temperature was maintained at 10 °C, and the injection volume was 1 µL. The chromatographic gradient was programmed as follows: 0–3 min, 3% B; 3–6 min, 3– 10% B; 6–21 min, 10–60% B; 21–23 min, 60–95% B; 23–26 min, 95% B; 26–27 min, 95–3% B; and 27–30 min, 3% B for column re-equilibration (Du et al. 2025). Aze and HAB were identified by matching retention times and accurate masses to those of authentic standards. Metabolite concentrations were determined using calibration curves generated from authentic Aze and HAB standards and normalized to tissue dry weight.

### RNA sequencing and transcriptomic analysis

WT and H-10 seedlings were grown for 8 d on half-strength Murashige and Skoog (½ MS) medium containing either 0 or 100 µM L-Aze. Whole seedlings were harvested, immediately frozen in liquid nitrogen, and stored at −80 °C until RNA extraction. Two independent biological replicates were collected for each genotype and treatment, resulting in four experimental groups: Col-0, Col-0 + 100 µM Aze, H-10, and H-10 + 100 µM Aze. Total RNA was isolated using the RNeasy Plant Mini Kit (QIAGEN) according to the manufacturer’s instructions. High throughput sequencing was performed at the University of Missouri Genomics Technology Core. Libraries were constructed following the manufacturer’s protocol with reagents supplied in Illumina’s Stranded RNA Prep, Ligation Kit (Illumina). Prior to library construction, the sample concentration were determined by Qubit fluorometer using the Qubit HS RNA assay kit (Invitrogen), and RNA integrity was checked using the Fragment Analyzer automated electrophoresis system (Agilent). Briefly, mRNA was captured with oligo(dT) magnetic beads. Captured RNA was eluted, fragmented and then primed for cDNA synthesis. First strand complementary DNA (cDNA) was synthesized using hexamer-primed RNA fragments and reverse transcriptase. After completion of second-strand cDNA synthesis, pre-index anchors were ligated to the ends of cDNA. A subsequent PCR step was used to selectively amplify the anchor-ligated DNA fragments and add unique dual indexes and primer sequences for cluster generation. The final amplified cDNA constructs were purified by addition of Axyprep Mag PCR Clean-up beads (Fisher Scientific). The final construct of each purified library was evaluated using the Fragment Analyzer automated electrophoresis system (Agilent), quantified with the Qubit fluorometer using the Qubit HS dsDNA assay kit (Invitrogen), and diluted according to Illumina’s standard sequencing protocol for sequencing on the NovaSeq X. The quality of the raw sequencing reads was assessed using FastQC v0.11.9, and quality-control reports were compiled using MultiQC v1.13 (Ewels et al. 2016). Adapter sequences and low-quality bases were removed using fastp v0.23.2. The quality of the filtered reads was reassessed before downstream analysis. Filtered reads were aligned to the *Arabidopsis thaliana* TAIR10 reference genome using HISAT2 v2.2.2 (Kim et al. 2019). Alignment files were converted to sorted BAM files using SAMtools v1.9 (Li et al. 2009) and gene-level read counts were generated using featureCounts from the Subread package with the corresponding TAIR10 gene annotation.

Differential gene expression was analyzed in R using DESeq2 (Love et al. 2014). Statistical testing was performed on raw gene-level counts using the negative-binomial model implemented in DESeq2. Genes with a Benjamini–Hochberg-adjusted *P* value below 0.01 and an absolute log fold change greater than 1 were considered differentially expressed. Variance-stabilizing transformation (VST) was applied to the count data for principal component analysis, sample-correlation analysis, hierarchical clustering, and heatmap visualization. VST-transformed values were used only for visualization and sample-level quality assessment and not for differential-expression testing. Heatmaps were generated in R using the pheatmap and ComplexHeatmap packages (Gu et al. 2016). For each selected gene set, VST expression values were standardized across samples by row-wise Z-score transformation. Both genes and samples were hierarchically clustered using Euclidean distance and Ward’s minimum-variance linkage method. Gene Ontology (GO) and Kyoto Encyclopedia of Genes and Genomes (KEGG) pathway-enrichment analyses were performed using gprofiler2 v0.2.1 (Raudvere et al. 2019). All genes included in the DESeq2 analysis were used as the statistical background. GO terms and KEGG pathways with a multiple-testing-adjusted (P) value below 0.05 were considered significantly enriched.

### Misincorporation proteomics analysis

WT and H-10 seedlings were grown for 8 d on half-strength Murashige and Skoog (½ MS) medium containing 100 µM L-Aze. Whole seedlings were harvested, immediately frozen in liquid nitrogen, and stored at −80 °C until protein extraction. Three pooled independent biological replicates were collected for each genotype. Proteins were extracted in 100 mM Tris pH 7.8, 10 mM DTT and 5% SDS (w/v). The protein was then heated at 65°C for 20 minute and centrifuged at 16,000 *× g* for 20 minutes. Proteins were alkylated with 30 mM iodoacetamide, precipitated using methanol/chloroform and washed once with 80% cold acetone. 50 μg of protein were then digested with LyC at 1:50 (enzyme:protein) for 3hrs at 37 °C and then digested with trypsin (1:50, enzyme:protein ratio) overnight at 37 °C. Digested peptides were purified by Evosep tips. Data were acquired on a Bruker timsTOF Pro2 connected to the Evosep-One system (Thives Santos et al., 2024). PEAKS version 10.6 was used to search the data against Arabidopsis TAIR10 database. Reversed protein sequences were appended to the original databases as decoys. For data analysis, precursor and fragment mass tolerances were set to 20 ppm, and 0.1 Da. Up to two missed trypsin cleavages were allowed. Oxidation of methionine, Pro to Aze (delta mass -14.0156 Da) were set as variable modifications. Carbamidomethylation of cysteine was set as a fixed modification. Maximum number of variable modifications per peptide was set to 3. Data were exported from PEAKS and peptides and proteins were filtered to FDR ≤0.01, Ascore ≥20. The protein quantification results from PEAKS Studio were exported and analyzed using Perseus 1.6.15.0. Proteins with at least one unique peptide in all three replicates and with valid peak area value in 70% of all samples were included for quantification.

### Statistical analysis

Statistical analyses were performed in R using RStudio. The experimental unit, sample size, and number of independent biological replicates are specified in the corresponding Figure legends or methods section. Comparisons between two independent groups were performed using a two-tailed Welch’s *t*-test when the assumptions for parametric analysis were satisfied. Experiments involving both genotype and treatment were analyzed by two-way analysis of variance (ANOVA), followed by Tukey’s HSD post hoc test. When parametric assumptions were not satisfied, the corresponding nonparametric tests were used. Concentration–response relationships and enzyme-kinetic parameters were estimated by nonlinear regression. Exact *P* values are reported whenever possible, and values below 0.001 are reported as *P* < 0.001. Differences were considered statistically significant at *P* < 0.05.

## Supplementary data

**Table S1.** Primers used in the study.

**Fig. S1.** Genotyping and molecular characterization of HAS-expressing transgenic lines.

**Fig. S2.** Phenotypic screening identifies H-10 as an Aze-tolerant HAS-expressing line.

**Fig. S3.** HAS expression enhances survival and reduces Aze sensitivity in H-10 plants.

**Fig. S4.** Phenotypic characterization of HAS-expressing H-10 plants under non-stress conditions.

**Fig. S5.** Computational prediction and experimental validation of the subcellular localization of HAS.

**Fig. S6.** Purification and kinetic characterization of recombinant HAS.

**Fig. S7.** Principal component and sample-correlation analyses reveal distinct transcriptomic responses to Aze in WT and H-10.

**Fig. S8.** Gene-set enrichment analysis reveals preservation of photosynthetic functions in H-10 and activation of proteotoxic-stress pathways in WT during Aze exposure.

**Fig. S9.** Aze induces phenylalanine, phenylpropanoid, and anthocyanin pathway genes more strongly in WT than in H-10.

**Fig. S10.** ER stress and protein quality-control responses are strongly induced in WT but attenuated in H-10 during Aze exposure.

**Fig. S11.** Unfolded protein response and protein-quality-control proteins accumulate to higher levels in Aze-treated WT than in H-10 plants.

**Fig. S12.** Potential active site residues involved in HAS reaction.

**Fig. S13.** Nucleotide sequence of the HAS coding region.

## Acknowledgements

We thank Brian Mooney and Thao Nguyen of the Charles W. Gehrke Proteomics Center at the University of Missouri for their assistance with the proteomics analyses. We are grateful to the Advanced Light Microscopy Core (AMLC) at MU for assistance with confocal microscopy. We also thank the Genomics Technology Core (RRID :SCR_017778) at the University of Missouri for providing RNA sequencing services. We thank the Waters Corporation (Milford, MA) for providing access to the Xevo MRT used in this study.

## Author contributions

VD, CAS conducted experiments, analyzed data, wrote and edited the manuscript. MB conducted experiments, analyzed data, and edited the manuscript.

## Funding

We acknowledge support from an MU College of Agriculture and Natural Resources Joy of Discovery grant for partial funding of this work.

## Data availability

The MS proteomics data have been deposited to the ProteomeXchange Consortium via the PRIDE partner repository with the dataset identifier PXD084193. RNAseq raw reads have been deposited to the sequence read archive (SRA) project number PRJNA1529264

## CONFLICT OF INTEREST STATEMENT

The authors declare no conflicts of interest.

## Notes

### Competing Interest Statement

The authors have declared no competing interest.

