## Supplementary Figures for "Engineering plants tolerant to the toxic proline mimic azetidine-2-carboxylic acid through co-option of a bacterial detoxification mechanism"

A

B

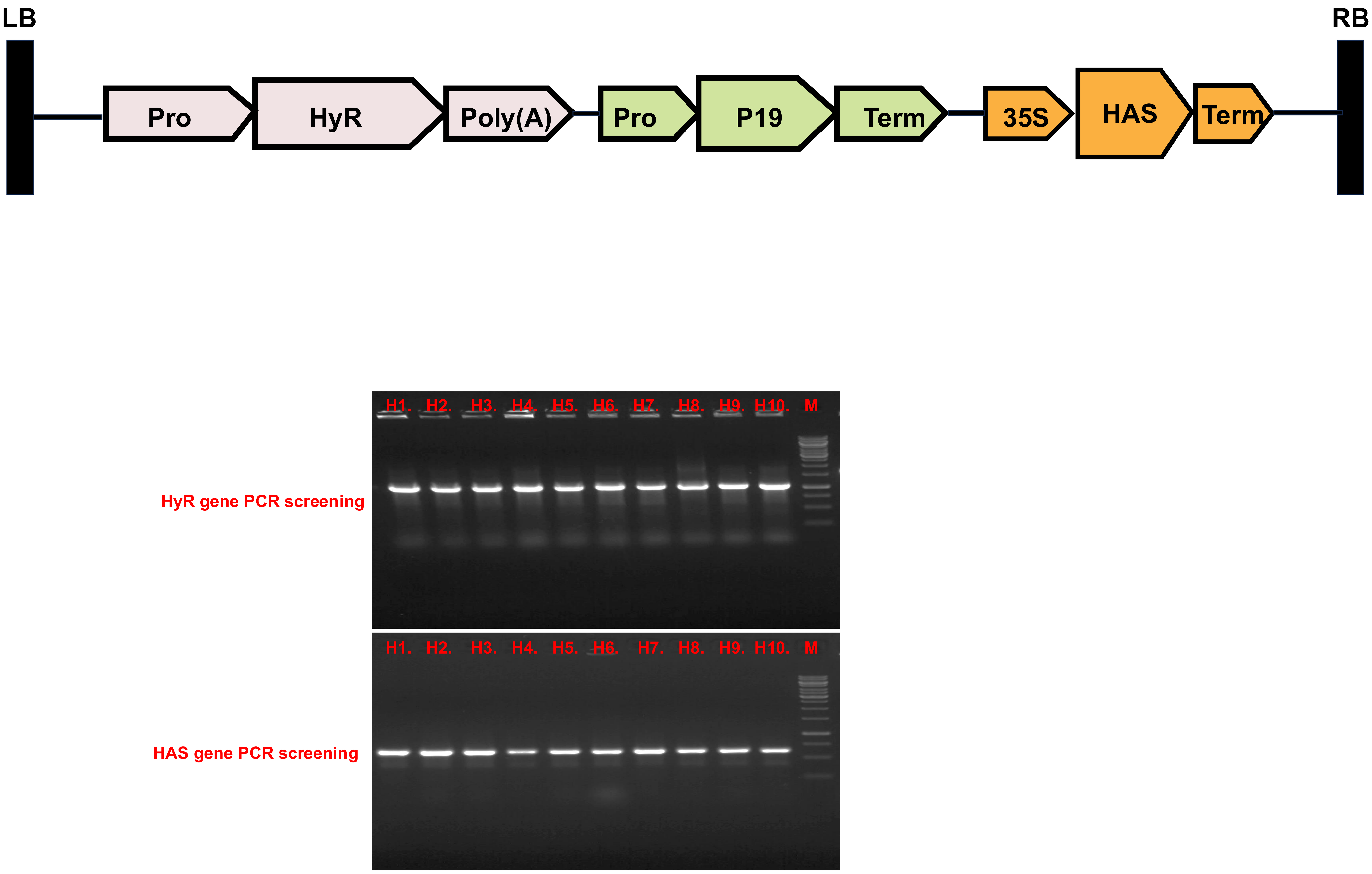


**Fig. S1. Genotyping of HAS-transgenic T0 lines.** (A) Schematic representation of the T-DNA construct used for constitutive expression of the bacterial Aze hydrolase HAS in *Arabidopsis thaliana*. The construct contains a hygromycin-resistance selectable-marker cassette (HyR), a P19 gene-silencing suppressor cassette, and a CaMV 35S-driven HAS expression cassette. Promoter (Pro), polyadenylation [poly(A)], and transcriptional terminator (Term) elements are indicated. The expression cassettes are flanked by the left and right T-DNA borders. The schematic is not drawn to scale. (B) PCR-based screening of 10 independent T0 transformation events (H1–H-10). The upper gel shows amplification of the HyR selectable-marker gene, and the lower gel shows amplification of the HAS transgene. PCR products of the expected sizes were detected in all 10 lines, confirming the presence of both transgenes. M, DNA size marker.

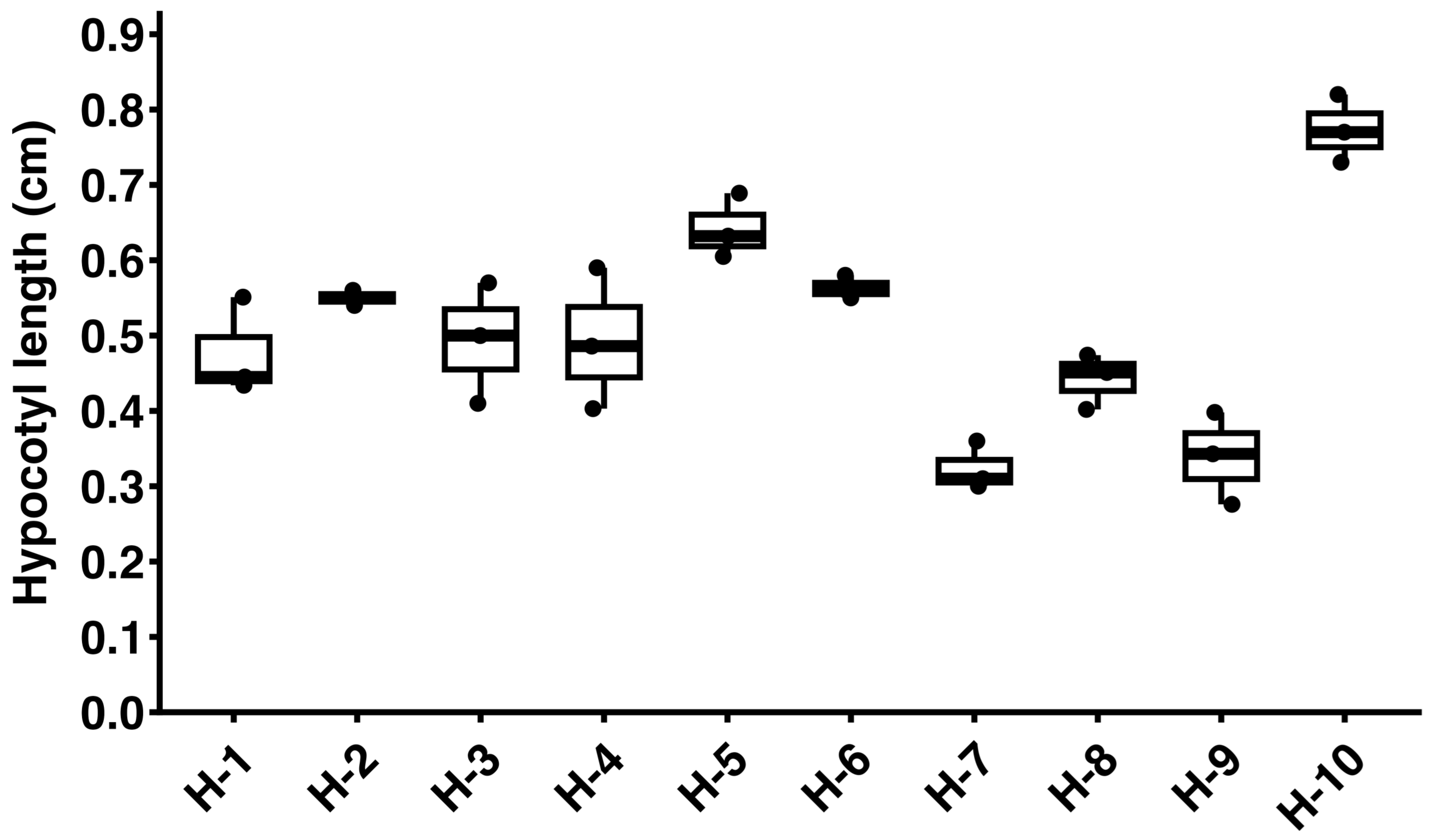


**Fig. S2. Phenotypic screening identifies H-10 as an Aze-tolerant HAS-expressing line.**
Seeds from independent HAS-expressing transgenic lines were germinated on ½ Murashige and Skoog (MS) medium supplemented with 250 μM azetidine-2-carboxylic acid (Aze). Hypocotyl length was measured as an indicator of Aze tolerance after 8 days of growth and used to distinguish strongly tolerant lines from lines exhibiting weaker tolerance. Line H-10 displayed the longest hypocotyl and was selected for subsequent physiological, biochemical, and molecular analyses.

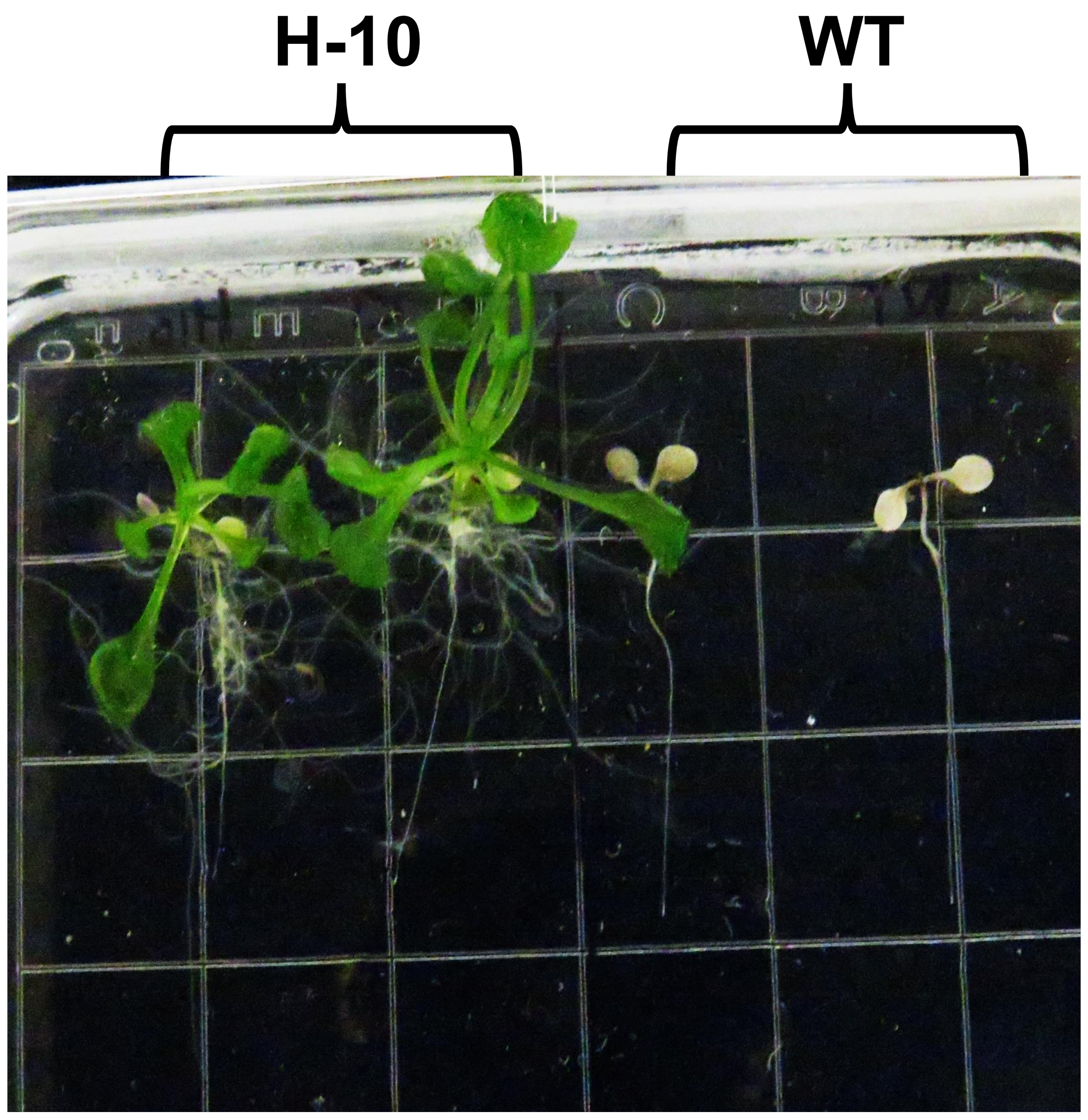


**Fig. S3. HAS expression enhances survival and reduces Aze sensitivity in H-10 plants.**
Representative image of WT and H-10 plant survival after growth on medium containing 500 μM Aze for 8 d. Quantified data found in Fig. 1D.

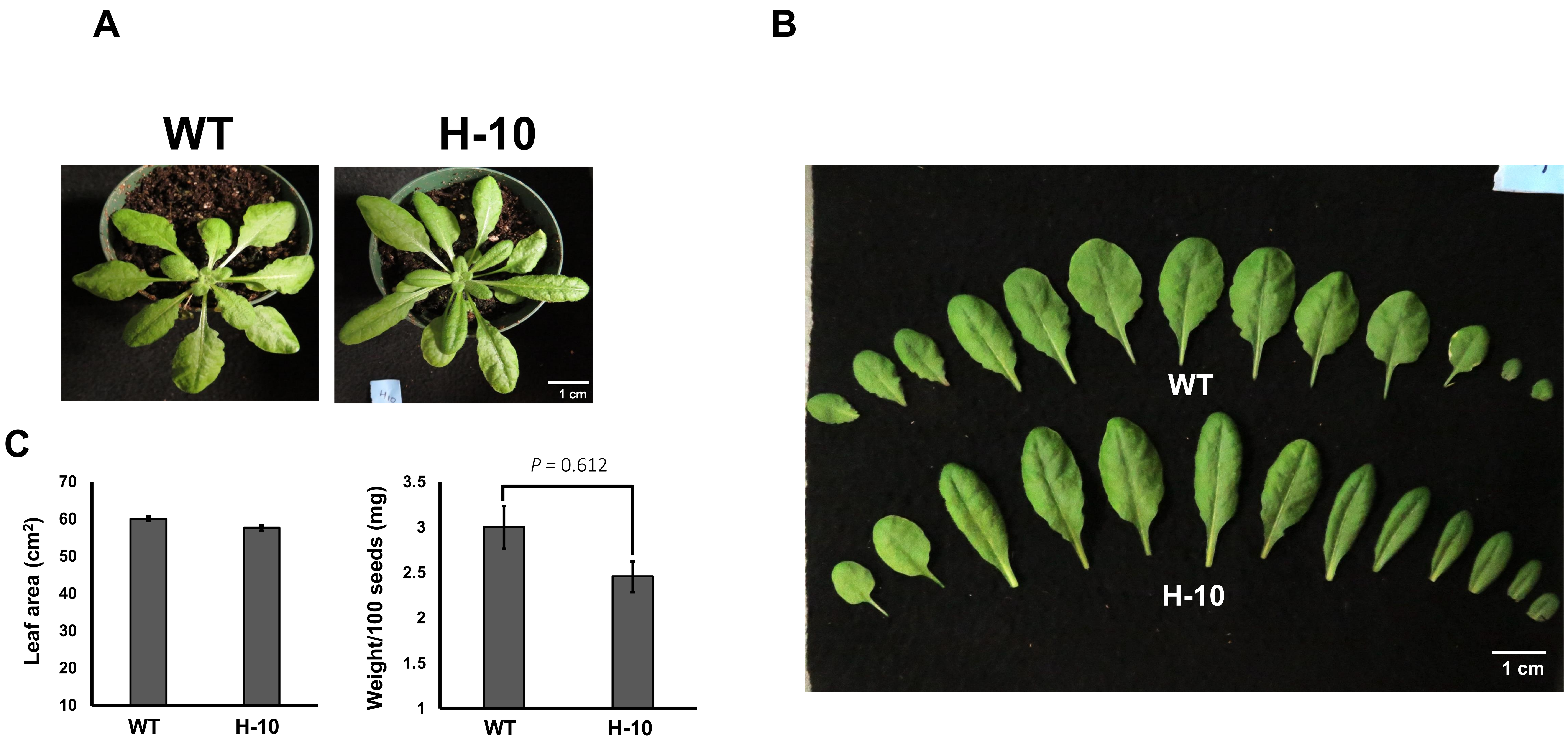


**Fig. S4. Phenotypic characterization of HAS-expressing H-10 plants grown in soil without Aze.**
(A) Representative rosette images of WT and H-10 plants grown under control conditions in the absence of azetidine-2-carboxylic acid (Aze). Scale bar, 1 cm. Images taken when plants were 4-5 weeks old.
(B) Representative images of detached rosette leaves from WT and H-10 plants, arranged according to developmental position. Scale bar, 1 cm.
(C) Quantification of total leaf area and the weight of 100 seeds from WT and H-10 plants. Leaf area was measured using ImageJ. Bars represent the mean ± SE from $n = 20$ independent biological replicates.

**A**

| Deeploc-2.0 | Cytoplasm | Nucleus | Extracellular | Cell membrane | Mitochondrion | Plastid | ER | Vacuole | Golgi apparatus | Peroxisome |
|---|---|---|---|---|---|---|---|---|---|---|
| HAS | 0.7001 | 0.3905 | 0.1058 | 0.2026 | 0.2707 | 0.0140 | 0.0780 | 0.1218 | 0.1581 | 0.0151 |

| Target P | Other | Signal peptide | Mitochondrial transfer peptide | Chloroplast transfer peptide | Thylakoid luminal transfer peptide |
|---|---|---|---|---|---|
| HAS | 0.9996 | 0.0002 | 0.0001 | 0.0 | 0.0 |

| WoLF PSORT | Cyto | Plas | Nuc | Cysk | Chlo | Mito | Pero |
|---|---|---|---|---|---|---|---|
| HAS | 3.5 | 2 | nucl: 5 | 3 | - | - | - |

| Predotar | Mitochondrial | Plastid | ER | Elsewhere | Prediction |
|---|---|---|---|---|---|
| HAS | 0,01 | 0,00 | 0,00 | 0,99 | none |

**B**

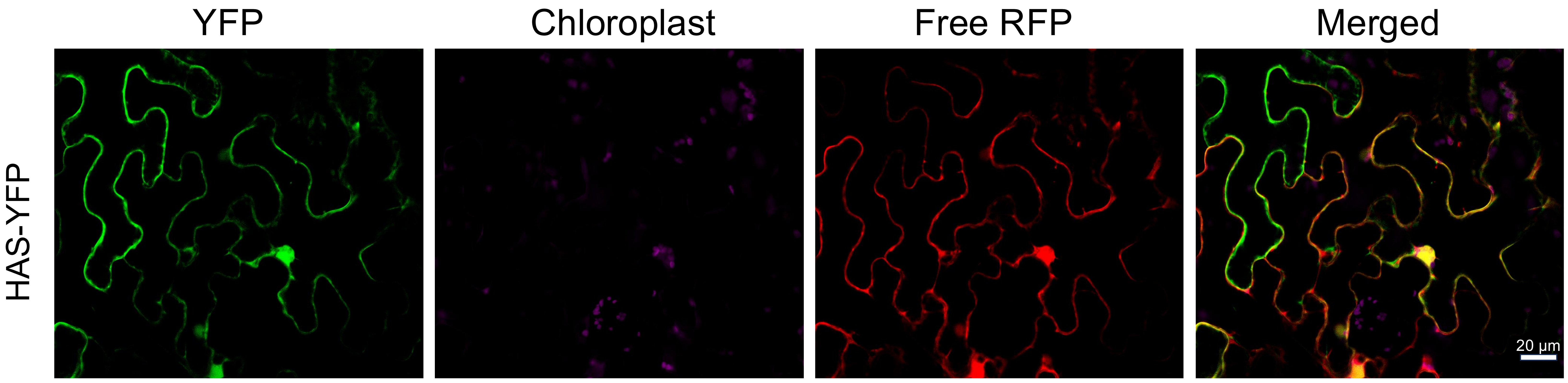


**Fig. S5. Computational prediction and experimental validation of the subcellular localization of HAS.**
(A) Subcellular localization of HAS predicted using DeepLoc-2.0, TargetP-2.0, WoLF PSORT, and Predotar. Values represent tool-specific prediction scores, with higher scores indicating stronger support for the corresponding localization within each platform. Collectively, the predictions indicated an absence of canonical plastidial, mitochondrial, or secretory targeting signal and supported a predominantly cytosolic localization.
(B) Representative confocal images of *Nicotiana benthamiana* epidermal cells transiently expressing HAS–YFP together with free red fluorescent protein (RFP) as a cytosolic marker. HAS–YFP fluorescence is shown in green, chlorophyll autofluorescence in magenta, and RFP fluorescence in red. The merged image shows colocalization of HAS–YFP with the cytosolic marker. Scale bar, 20 µm.

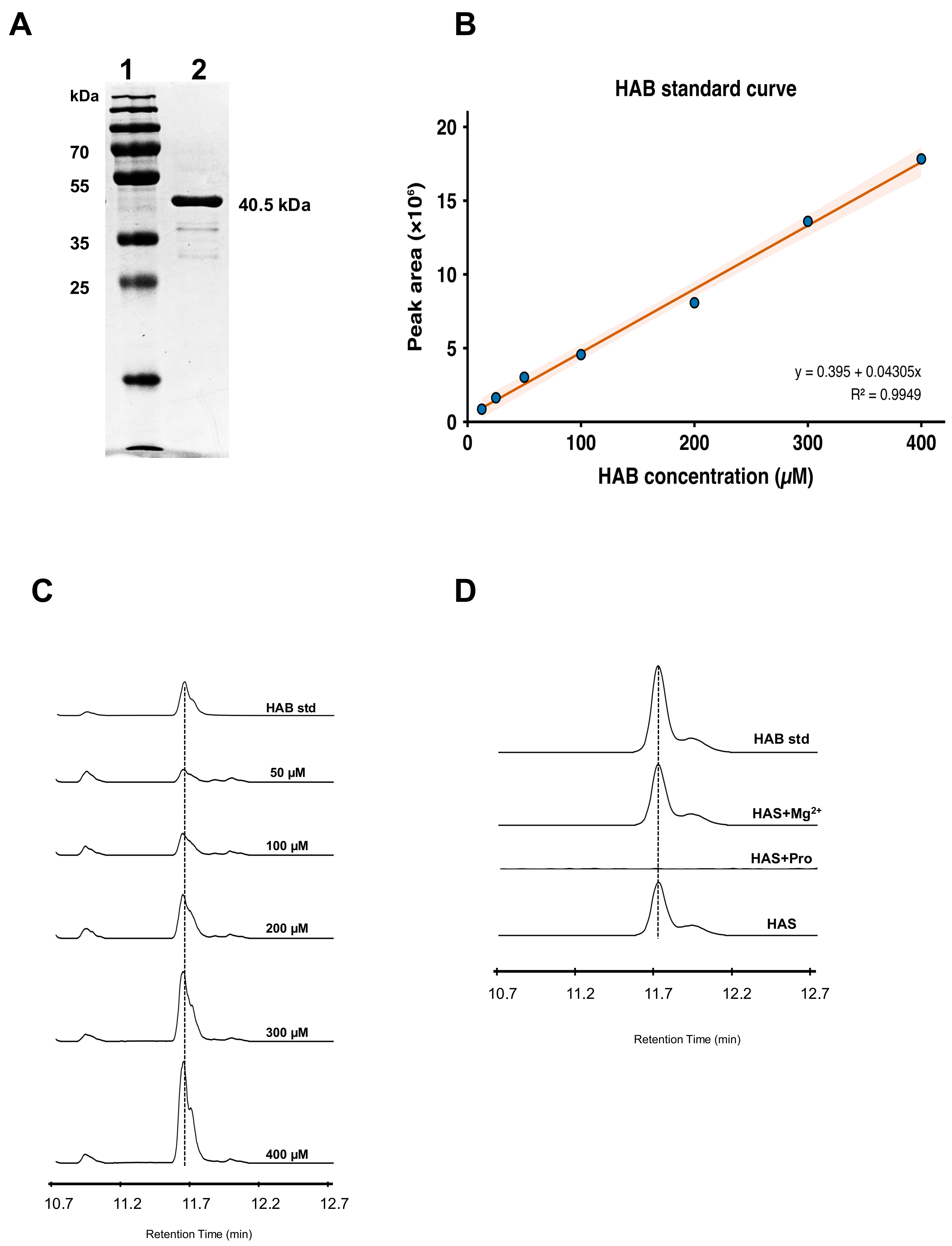


**Fig. S6. Purification and *in vitro* enzymatic characterization of recombinant HAS.**
(A) Coomassie-stained SDS–polyacrylamide gel showing nickel-affinity-purified recombinant His-tagged HAS expressed in *Escherichia coli*. Lanes 1, molecular-weight marker; lane 2, purified HAS used for enzyme assays. **(B)** HAB standard curve generated by plotting LC–MS peak area against HAB concentration (12.5–400 μM); the line represents the linear regression fit ($R^2$ = 0.99). **(C)** Representative LC–MS chromatograms showing HAB formation in reactions containing 0.25 μM purified HAS and increasing concentrations of L-Aze. Reactions were incubated for 5 min before termination. The upper trace represents the authentic HAB standard, and the dashed line indicates its retention time. **(D)** Representative LC–MS chromatograms showing HAB formation in reactions containing purified HAS, $Mg^{2+}$, and L-Aze. HAB was not detected when L-proline was used as a substrate. Product identity was confirmed by comparison of its retention time and intact mass and fragmentation pattern with that of the authentic HAB standard.

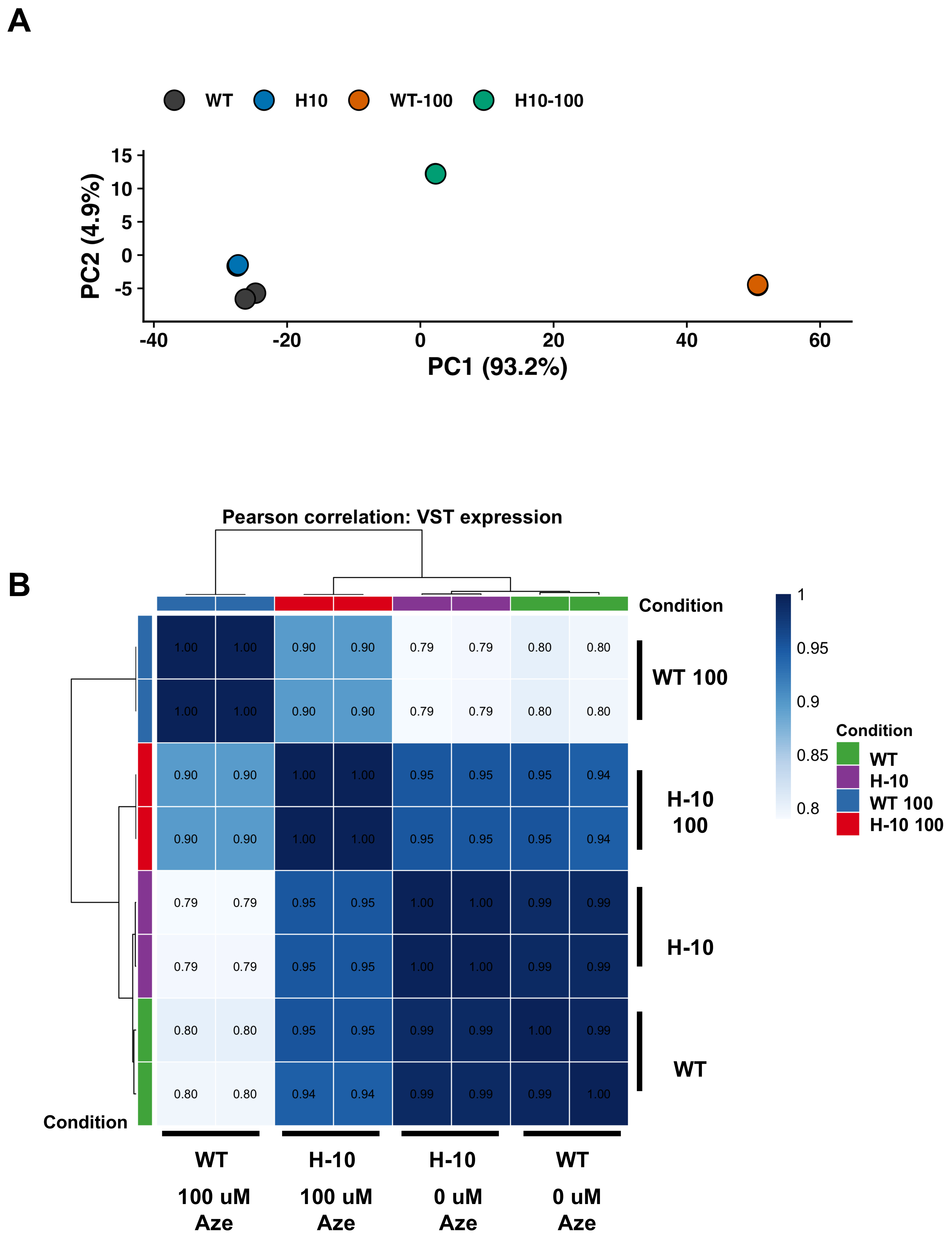


**Fig. S7. Principal component and sample-correlation analyses reveal distinct transcriptomic responses to Aze in WT and H-10.** (A) Principal component analysis of variance-stabilized RNA-seq expression data from untreated and 100 μM Aze-treated WT and H-10 seedlings. PC1 and PC2 accounted for 93.2% and 4.9% of the total variance, respectively. Each point represents an independent biological replicate, and colors indicate the corresponding genotype and treatment. Overlapping points reflect highly similar expression profiles between replicates. At a global level, WT and H-10 showed very similar transcriptional patterns without Aze treatment suggesting little transcriptional impact of HAS expression.

(B) Pearson correlation analysis of RNA-seq samples. Heatmap showing pairwise Pearson correlation coefficients calculated from variance-stabilized gene-expression values across all RNA-seq samples. Values within cells indicate the correlation coefficients, and dendrograms show hierarchical clustering based on global expression profiles. Two independent biological replicates were analyzed per condition. WT, untreated Col-0 wild type; H-10, untreated HAS-overexpressing line; WT-100 and H-10-100, the corresponding genotypes treated with 100 μM Aze.

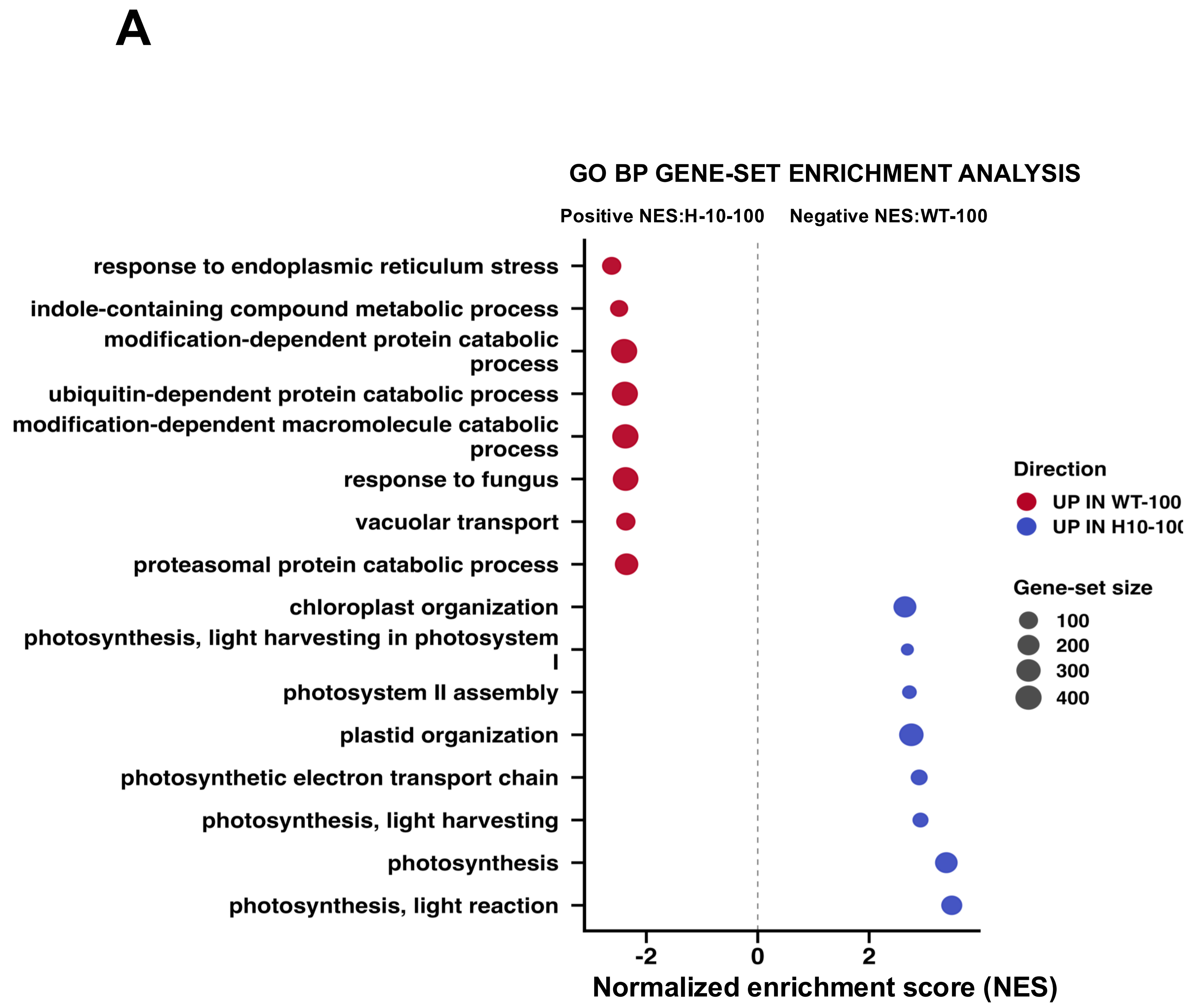


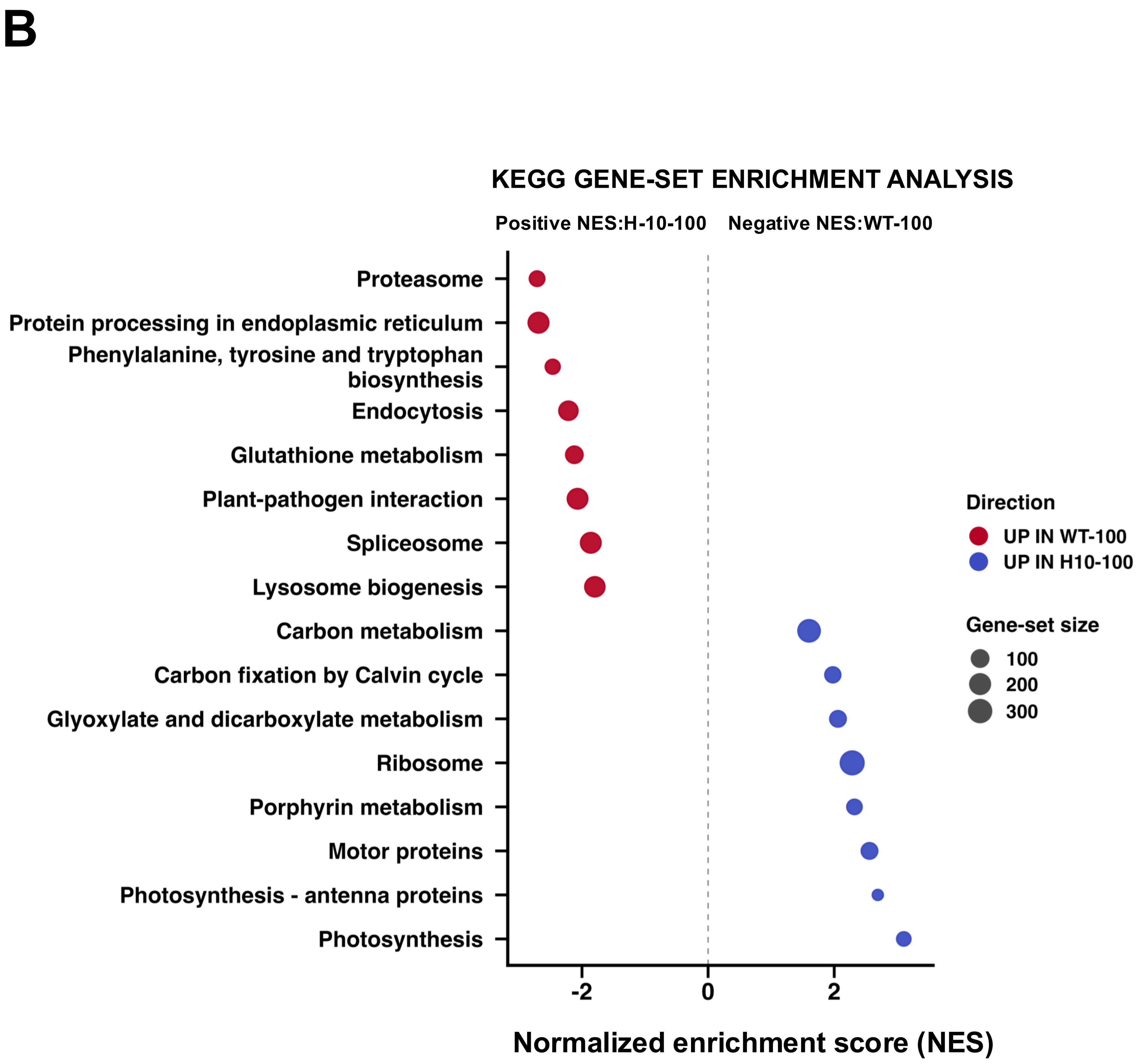


**Fig. S8. Gene-set enrichment analysis reveals preservation of photosynthetic functions in H-10 and activation of proteotoxic stress pathways in WT during Aze exposure.** Gene-set enrichment analysis (GSEA) was performed using the complete ranked gene list from the H-10-100 versus WT-100 comparison. The ranked list included all genes tested for differential expression, ordered by the differential-expression ranking statistic without applying significance or fold-change cutoffs. Positive normalized enrichment scores (NESs) indicate gene sets enriched among genes expressed more highly in H-10-100 (blue), whereas negative NESs indicate gene sets enriched among genes expressed more highly in WT-100 (red). In both panels, point position indicates the NES, point size represents gene-set size, and the dashed vertical line marks an NES of zero. (A) Gene Ontology Biological Process (GO BP) analysis. Photosynthesis, photosynthetic light reactions and light harvesting, photosystem II assembly, photosynthetic electron transport, and plastid and chloroplast organization were enriched in H-10-100. In contrast, responses to endoplasmic reticulum stress and fungi, indole-containing compound metabolism, modification- and ubiquitin-dependent protein catabolism, proteasomal protein catabolism, and vacuolar transport were enriched in WT-100. (B) Kyoto Encyclopedia of Genes and Genomes (KEGG) pathway analysis. Photosynthesis, photosynthesis–antenna proteins, porphyrin metabolism, ribosome, motor proteins, glyoxylate and dicarboxylate metabolism, carbon fixation by the Calvin cycle, and carbon metabolism were enriched in H-10-100. Protein processing in the endoplasmic reticulum, proteasome, phenylalanine, tyrosine, and tryptophan biosynthesis, endocytosis, glutathione metabolism, plant–pathogen interaction, spliceosome, and lysosome biogenesis were enriched in WT-100. H-10-100 and WT-100 denote H-10 and Col-0 wild-type seedlings, respectively, treated with 100 μM Aze.

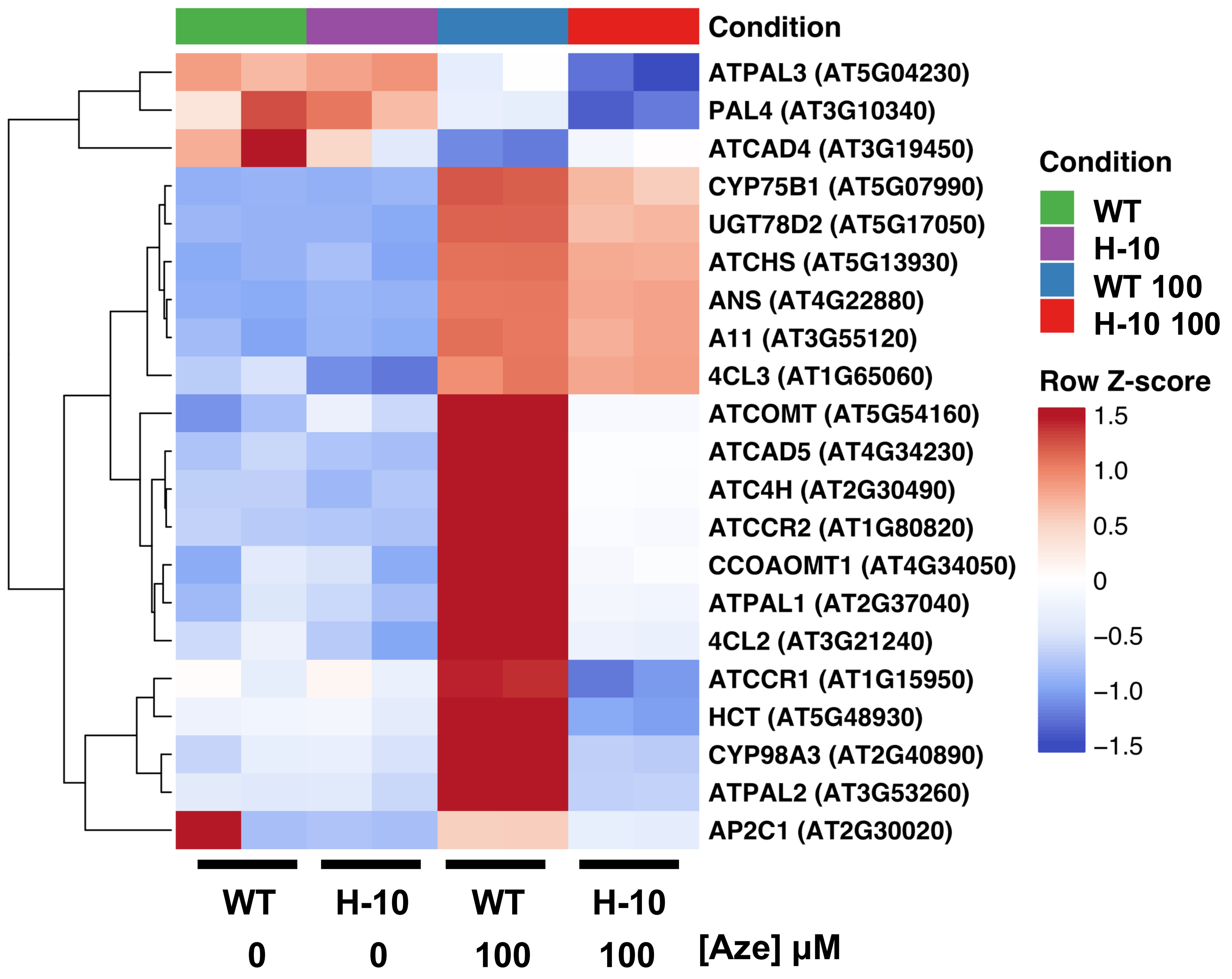


**Fig. S9. Aze induction of phenylalanine, phenylpropanoid, and anthocyanin pathway genes is repressed in H-10.**
Heatmap showing the expression of selected genes associated with phenylalanine, phenylpropanoid, and anthocyanin metabolism in untreated and 100 µM Aze-treated WT and H-10 seedlings. Variance-stabilized expression values were standardized independently for each gene using row-wise Z-score transformation. Red and blue indicate higher and lower relative expression, respectively, with the color scale limited to −1.5 to 1.5. Genes were hierarchically clustered based on their expression patterns. Each condition contains two independent biological replicates. Core pathway genes, including *PAL1*, *PAL2*, *C4H*, *4CL2*, *CCR1*, *HCT*, *CHS*, *ANS*, *CYP75B1*, and *UGT78D2*, were strongly induced in WT-100 but showed weaker induction in H-10-100. WT-100 and H-10-100 denote WT and H-10 seedlings, respectively, treated with 100 µM Aze.

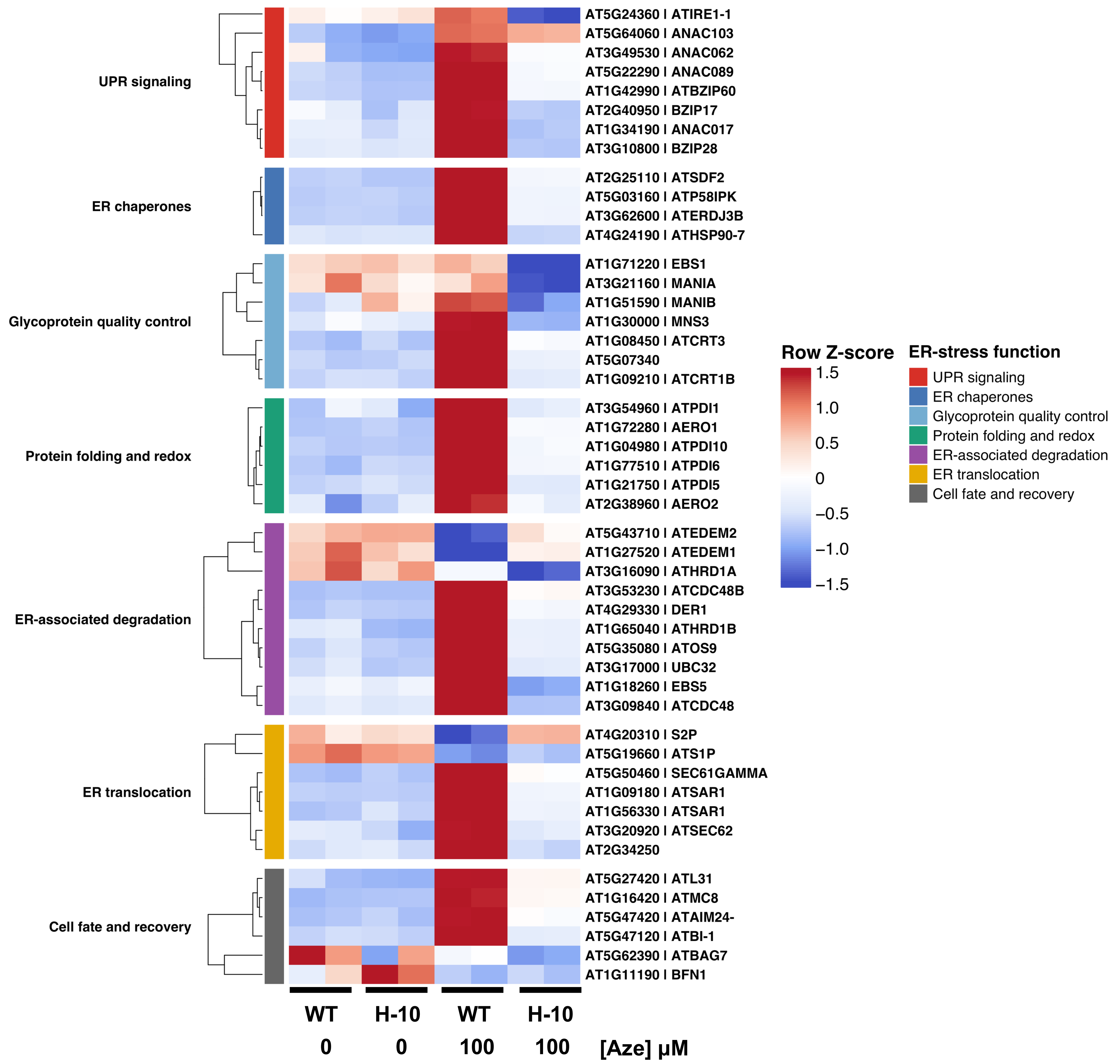


**Fig. S10. ER stress and protein quality-control responses are strongly induced in WT but attenuated in H-10 during Aze exposure.**

Heatmap showing the 48 most variable genes associated with endoplasmic reticulum stress and protein quality control in untreated and 100 μM Aze-treated WT and H-10 seedlings. Genes are organized into seven functional categories: unfolded protein response signaling, ER chaperones, glycoprotein quality control, protein folding and redox regulation, ER-associated degradation, ER translocation, and cell fate and recovery. Variance-stabilized expression values were standardized independently for each gene using row-wise Z-score transformation. Red and blue indicate higher and lower relative expression, respectively, with the scale limited to −1.5 to 1.5. Genes were hierarchically clustered within each functional category. Each condition contains two independent biological replicates. A broad set of ER stress-associated genes was strongly induced in WT-100, whereas most showed substantially weaker induction in H-10-100. WT-100 and H-10-100 denote WT and H-10 seedlings, respectively, treated with 100 μM Aze.

A

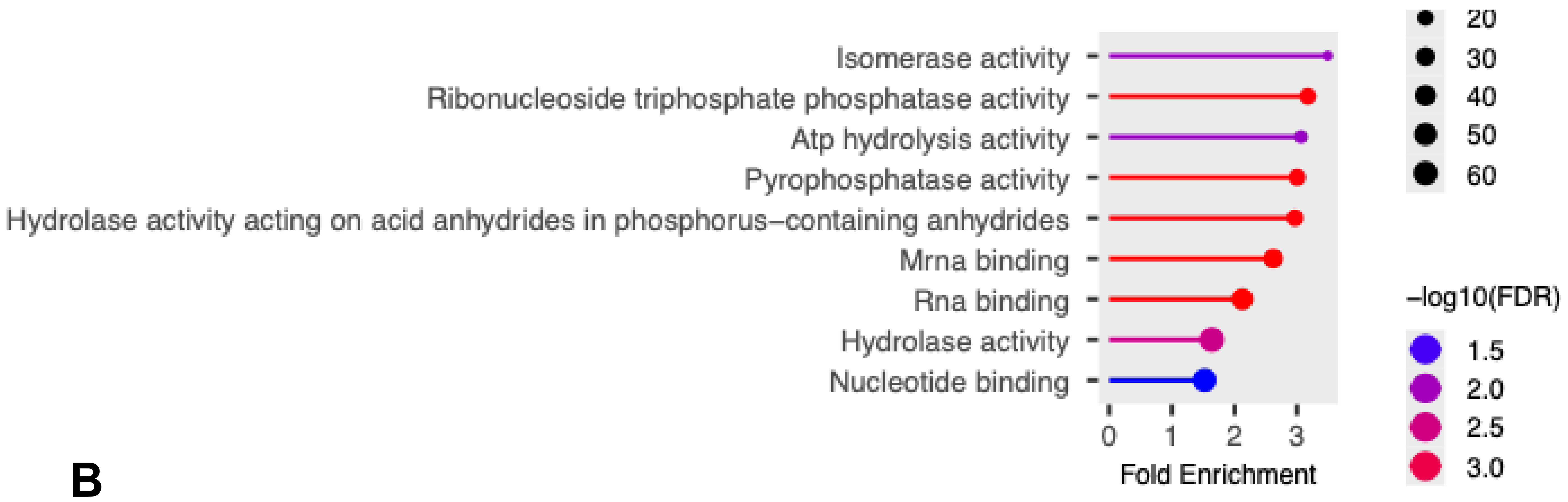


B

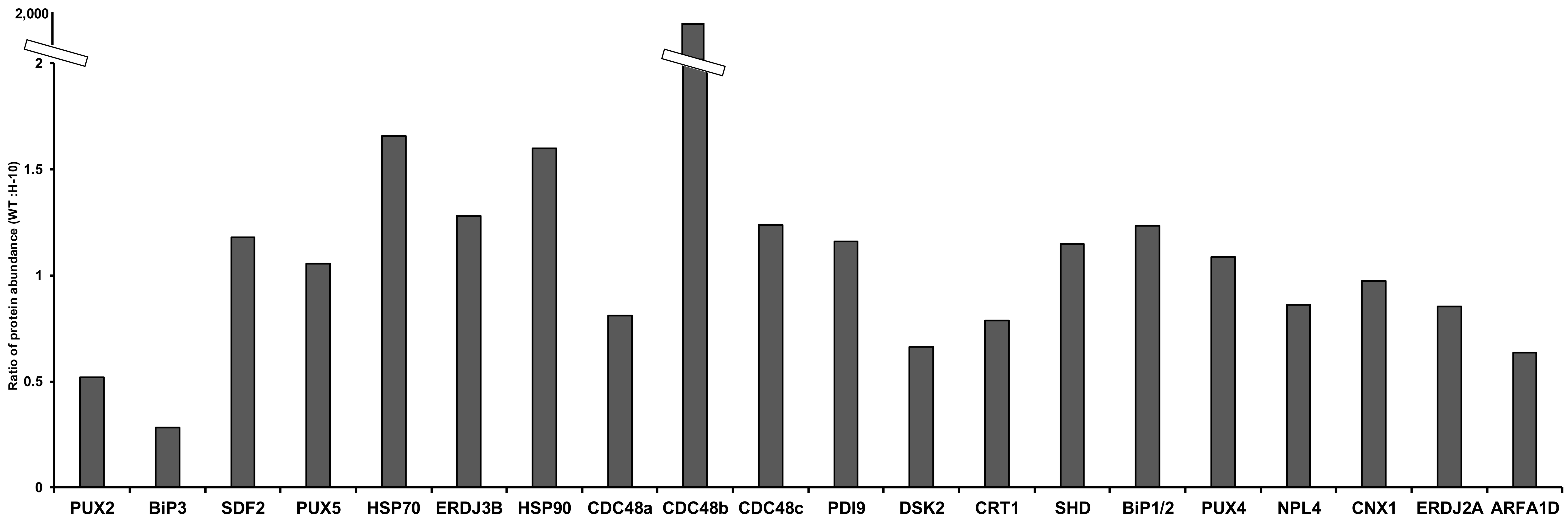


**Fig. S11. Unfolded protein response and protein-quality-control proteins accumulate to higher levels in Aze-treated WT than in H-10 plants.** (A) Functional enrichment analysis using proteins with a 2-fold increased abundance in WT compared to H-10 (321 total proteins). (B) Relative abundance of selected proteins associated with the unfolded protein response and protein quality control in Aze-treated WT and H-10 plants. A ratio of protein abundance in WT compared to H-10 during Aze treatment is shown. Values greater than 1 indicate higher protein abundance in WT.

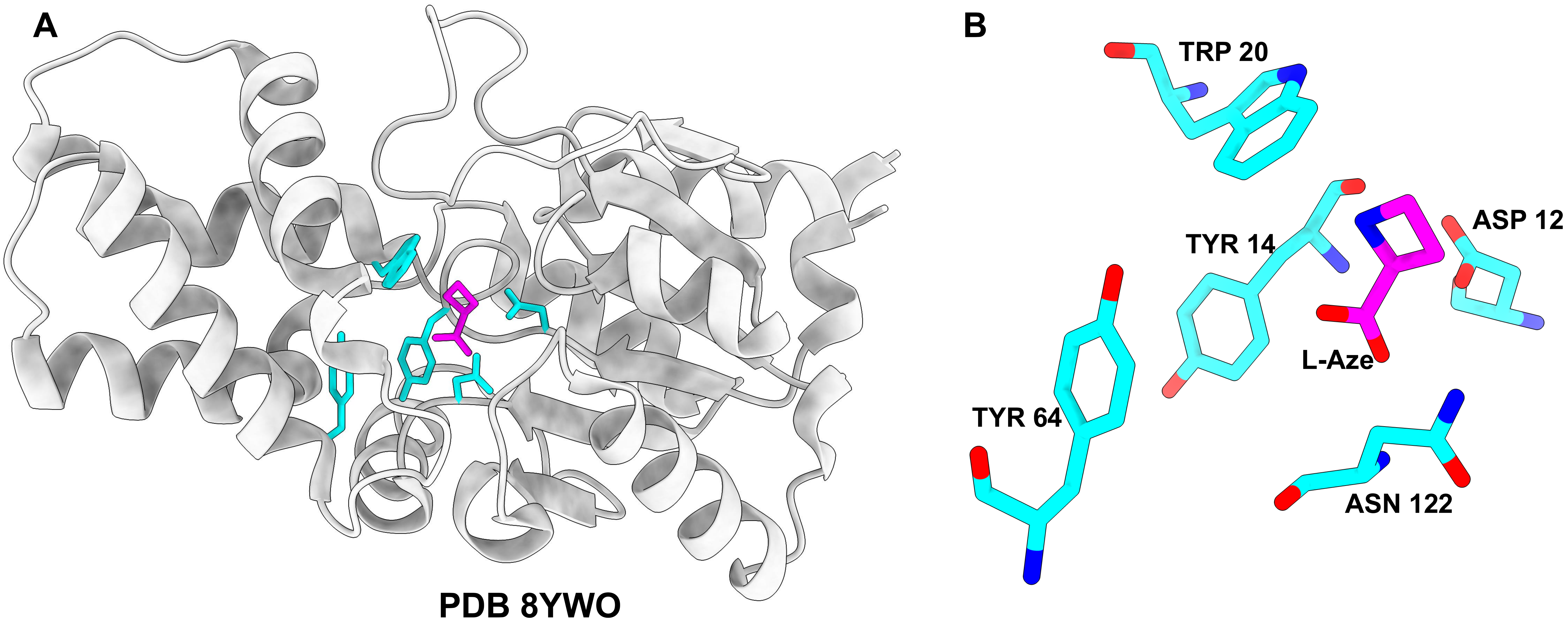


**Fig. S12. Potential active site residues involved in HAS reaction.**
**(A)** Overall structure of HAS in complex with L-azetidine-2-carboxylic acid (L-Aze) (PDB 8YWO). HAS is shown as a light-gray cartoon and L-Aze as magenta sticks. **(B)** Ligand-focused enlarged view of the predicted HAS active-site pocket showing L-Aze (magenta sticks), and the surrounding residues Asp12, Trp20, Tyr64, and Asn122 (Cyan sticks). The position of Asp12 near both L-Aze confirms its role in substrate binding or catalysis.

```
         10        20        30        40        50
ATGCAGCTCACCGACTTTAAAGCTCTTACCTTTGACTGCTATGGAACGCT    50
TATCGACTGGGAGACAGGCATTGTCAACGCTTTACAACCTTTGGCAAAAA    100
GGACGGGGAAAACATTCACATCTGACGAATTGCTGGAAGTTTTCGGACGT    150
AACGAATCTCCCCAGCAGACGGAAACTCCCGGAGCACTTTATCAGGATAT    200
TTTAAGGGCCGTTTACGACAGGATAGCCAAGGAGTGGGGATTAGAGCCAG    250
ACGCTGCGGAACGTGAGGAGTTCGGTACTAGTGTTAAAAACTGGCCGGCG    300
TTTCCAGATACCGTGGAAGCACTACAATACCTAAAAAAACACTATAAACT    350
GGTGATATTAAGCAACATAGATCGTAACGAGTTTAAGTTGAGCAATGCTA    400
AACTGGGAGTGGAGTTTGATCACATCATCACGGCACAGGATGTGGGAAGT    450
TACAAGCCTAATCCCAACAACTTCACTTACATGATAGATGCGCTTGCAAA    500
AGCAGGCATTGAAAAGAAGGACATTCTCCATACTGCTGAGTCACTTTATC    550
ACGACCATATTCCCGCCAATGACGCAGGCCTAGTCTCCGCTTGGATATAC    600
AGAAGACACGGCAAGGAGGGATACGGAGCAACCCACGTACCATCAAGGAT    650
GCCGAATGTCGATTTCCGATTCAACTCCATGGGTGAGATGGCAGAGGCAC    700
ATAAGCAAGCCTTGAAGGGCTAA                               723
```

**Fig. S13. Nucleotide sequence of the HAS coding region.** The complete 723-bp coding sequence of the bacterial L-azetidine-2-carboxylic acid hydrolase (*HAS*) is presented in the 5′-to-3′ direction. The sequence begins with the ATG start codon and terminates with the TAA stop codon, encoding a protein of 240 amino acids. Numbers on the right indicate nucleotide positions.
